# Retrospective attention strengthens a report-independent neural signature of visual perception

**DOI:** 10.1101/2024.11.09.622275

**Authors:** Ye Li, Xingbei Wan, Mingxing Mao, Lingying Jin, Yaochun Cai, Xilin Zhang

**Author notes:** Corresponding authors: Ye Li, Xilin Zhang, (YL); (XZ).

## Abstract

Visual perception can remain modifiable after a stimulus has disappeared, but it is unknown at which processing stage a post-stimulus cue operates: on the lingering sensory trace, on the perceptual representation, or on later decision and report processes. We used masked retrospective cues to establish the behavioral effect and a Report/No-report EEG design to locate its processing stage. In Experiments 1 and 2, retrospective cues improved orientation discrimination even when masking reduced cue-location accuracy to near chance. In Experiment 3, valid cues increased EEG evidence for the target in a late interval, without the early enhancement expected from an early, additive sensory interaction. Without explicit target report, valid cues still strengthened the neural representation of the target, and a decoder trained only on no-report trials predicted trial-by-trial visibility and orientation discrimination in separate report trials, arguing against a purely report-contingent decisional account. When report was required, the cueing benefit appeared mainly in additional report-specific signals, which also predicted visibility most strongly. Retrospective attention thus strengthens a report-independent visual target representation, and explicit report can amplify this effect.

## Introduction

Visual perceptual processing can remain modifiable even after a stimulus has disappeared. In pioneering work on retroperception, Sergent and colleagues showed that directing retrospective attention to a near-threshold Gabor patch after its offset significantly improved both orientation discrimination and subjective visibility[1], suggesting that retrospective attention can modulate conscious access to past sensory representations[2]. These findings show that the fate of a visual representation is not fixed at stimulus offset[1, 2]. However, it remains unresolved at which processing stage a post-stimulus cue operates: on the lingering sensory trace, on the perceptual representation itself, or on downstream decision, motor, or memory processes[3, 37]. Retrospective cueing is uniquely suited to this question, because the cue arrives after initial encoding and can therefore act only on what the stimulus has left behind.

The answer bears on a core debate about attention and consciousness: whether the two operate independently[5–7] or whether attentional selection is a gateway to conscious access[8, 9]. Three accounts make distinct predictions about where the cue operates (Fig 1; Table 1). A sensory-interaction account holds that the cue interacts locally with the lingering sensory trace at the cued location[23, 41–43]; it predicts that valid cues raise target evidence early, within the window of sensory encoding, irrespective of report. A decisional account holds that the cue acts only on post-perceptual decision, working-memory, or response processes[3, 10–12, 37, 44, 45]; it predicts no cueing benefit without report, and neural activity recorded without report should not predict perception. A representational account holds that the cue strengthens a target representation that is still available after the stimulus is gone, so that it can still reach conscious access[1, 2, 16, 38, 40]; it predicts a late increase in target evidence even without report, and this evidence should predict visibility and discrimination. These accounts describe limiting cases rather than mutually exclusive mechanisms; the question is which stage carries the cueing effect. Standard paradigms cannot separate these accounts for two reasons. First, when participants report the target, post-perceptual processes can hide or imitate changes in the perceptual representation[3, 10–12]. Second, clearly visible cues allow deliberate, strategic orienting; masking the cue tests whether spatial selection still occurs when participants can barely locate the cue[4, 13, 14].

**Fig 1.**
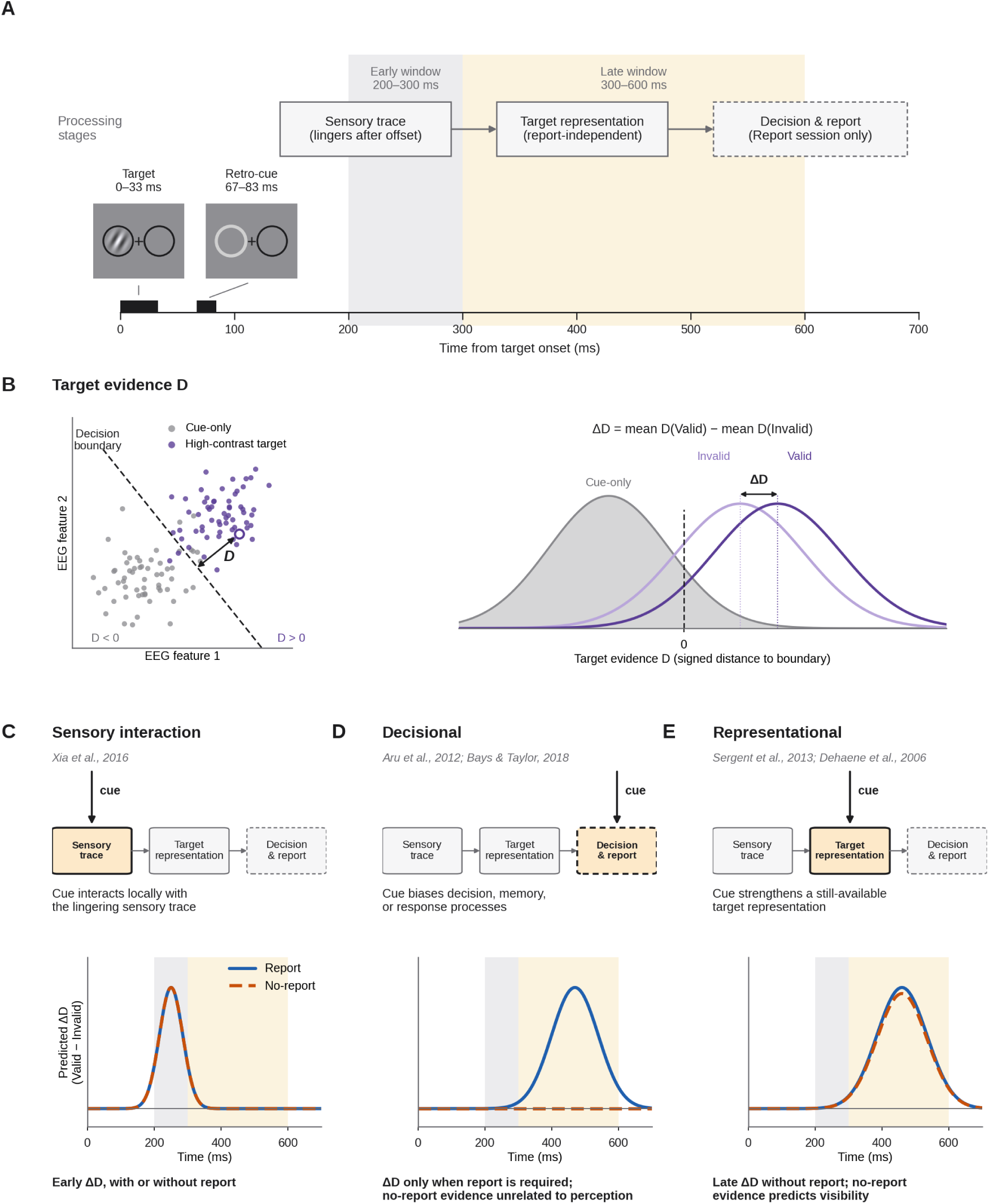
Three accounts of where a retrospective cue operates, and their predictions. (A) Trial structure and processing stages. A near-threshold Gabor target (0–33 ms) is followed by a peripheral retrospective cue (67–83 ms). After stimulus offset, the target leaves a lingering sensory trace, which can give rise to a report-independent target representation. When a report is required, decision and report processes follow (dashed box). Shading marks the early (200–300 ms) and late (300–600 ms) analysis windows. (B) Target evidence D. A decoder trained to separate High-contrast target trials from cue-only trials defines a decision boundary in EEG feature space; D is the signed distance of each held-out trial from this boundary (positive on the target side). The cueing effect is ΔD, the mean D of Valid minus Invalid trials. (C–E) Each account places the action of the cue at a different stage (highlighted box) and predicts a different time course of the cueing effect on target evidence (ΔD, Valid − Invalid) in the Report (solid line) and No-report (dashed line) sessions. (C) A sensory-interaction account predicts an early effect, with or without report. (D) A decisional account predicts an effect only when report is required, and no link between neural evidence recorded without report and perception. (E) A representational account predicts a late effect even without report, carried by a representation that predicts visibility. Panels B–E are schematic, not data.

**Table 1.** Competing accounts of retrospective cueing, their predictions, and the corresponding results.

| Account | Proposed locus of the cue effect | Prediction | Test | Result |
| --- | --- | --- | --- | --- |
| Sensory interaction | Lingering sensory trace at the cued location[23, 41–43] | Cueing effect within the window of sensory encoding (200–300 ms), irrespective of report | Time course of target evidence (Experiment 3) | No early enhancement (early posterior effects reversed); corrected enhancement after 300 ms (S9 and S12 Figs) |
| Decisional | Post-perceptual decision, memory, or response processes[3, 37, 44, 45] | Report-contingent form: no cueing effect when report is not required; evidence acquired without report unrelated to perception | Cueing under No-report; No-report-trained decoder tested on Report behavior (Experiment 3) | Cueing effect without report; No-report representation predicts Seen responses and accuracy (Fig 4) |
| Representational | Still-available target representation[1, 2, 38] | Late enhancement of report-independent target evidence that predicts perception | Decomposition into shared and Report-specific components (Experiment 3) | Shared representation enhanced without report; report amplifies the effect in report-specific signals when the Report session comes first (Fig 5E–F) |

No-report paradigms separate perceptual representation from report-related activity[3, 5]. Recent studies show that some neural markers of conscious perception persist without explicit reports, whereas classic late signals depend strongly on whether a report is required[15–19]. In audition, late activity beyond approximately 250 ms persisted during passive listening and predicted what participants reported at intermittent probes[16]. In visual masking, fronto-central activity around 250–300 ms tracked perceptual awareness across report and no-report conditions, whereas the late P3b and decoding that remained stable over time were tied mainly to explicit report[17]. Cross-task decoding has extended this logic: a ventral visual signal that tracked detection reports also distinguished stimulus intensity during passive viewing[20], and information about conscious content generalized across task relevance, most robustly in posterior cortex[21]. However, these studies manipulated stimulus visibility, strength or task relevance, not retrospective attention. It is therefore unknown whether a post-stimulus cue can strengthen a target representation when no report is required, and whether such a representation is linked to conscious perception[2, 22, 23].

Here, in three experiments, we tested these accounts by masking the cue (Experiments 1 and 2) and by removing explicit target report (Experiment 3). In Experiments 1 and 2, masked retrospective cues improved orientation discrimination even though participants could barely tell where the cue had appeared. In Experiment 3, the same participants completed Report and No-report sessions in counterbalanced order while we recorded EEG; we subtracted cue-evoked activity and used a decoder trained without regard to cue validity[26–28]. Each account was put to a direct test. If the cue interacts with the lingering sensory trace, the cueing effect should appear early, during sensory encoding. If the cue acts only through report, the effect should disappear without report, and a decoder trained without report should not predict visibility or orientation discrimination in the Report session. If the cue strengthens the target representation itself, the effect should appear without report, on a representation shared by both sessions. We find that retrospective attention strengthens a report-independent target representation that predicts visibility and discrimination, and that explicit report can amplify this effect through additional report-specific signals.

## Results

### Retrospective cues improve discrimination under strongly reduced cue information

Experiments 1 and 2 first established the behavioral benefit of retrospective cueing at the interval used throughout. In Experiment 1 (N = 24), participants discriminated the orientation of a near-threshold Gabor target presented left or right of fixation; a peripheral dimming cue appeared at the target location (Valid) or the opposite location (Invalid) before (−116.67 ms) or after the target (+66.67 or +100 ms), was either visible or immediately backward-masked, and was followed by a cue-location 2AFC that measured residual cue information (Fig 2A). Orientation discrimination was reliably higher after Valid than Invalid cues (trial-level GLMM, F(1,23124) = 48.27, Holm-adjusted p = 1.53 × 10⁻¹¹); the benefit was modulated by cue visibility (Validity × Cue visibility, Holm-adjusted p = .0013) but did not differ between pre- and post-target cues (Validity × SOA, Holm-adjusted p = .440). With masked cues, orientation accuracy at the +66.67 ms interval was higher for Valid than Invalid trials by 3.04 percentage points (t(22) = 2.20, p = .038; Fig 2D; this cell-level difference depended on the 65–85% inclusion criterion, whereas the trial-level validity effect held when all 29 recruited participants were included; S1 Table), while cue-location accuracy remained within ±5 percentage points of chance (52.69% at +66.67 ms; 50.17% overall; Fig 2B; S1 Fig; S1 Table).1

**Fig 2.**
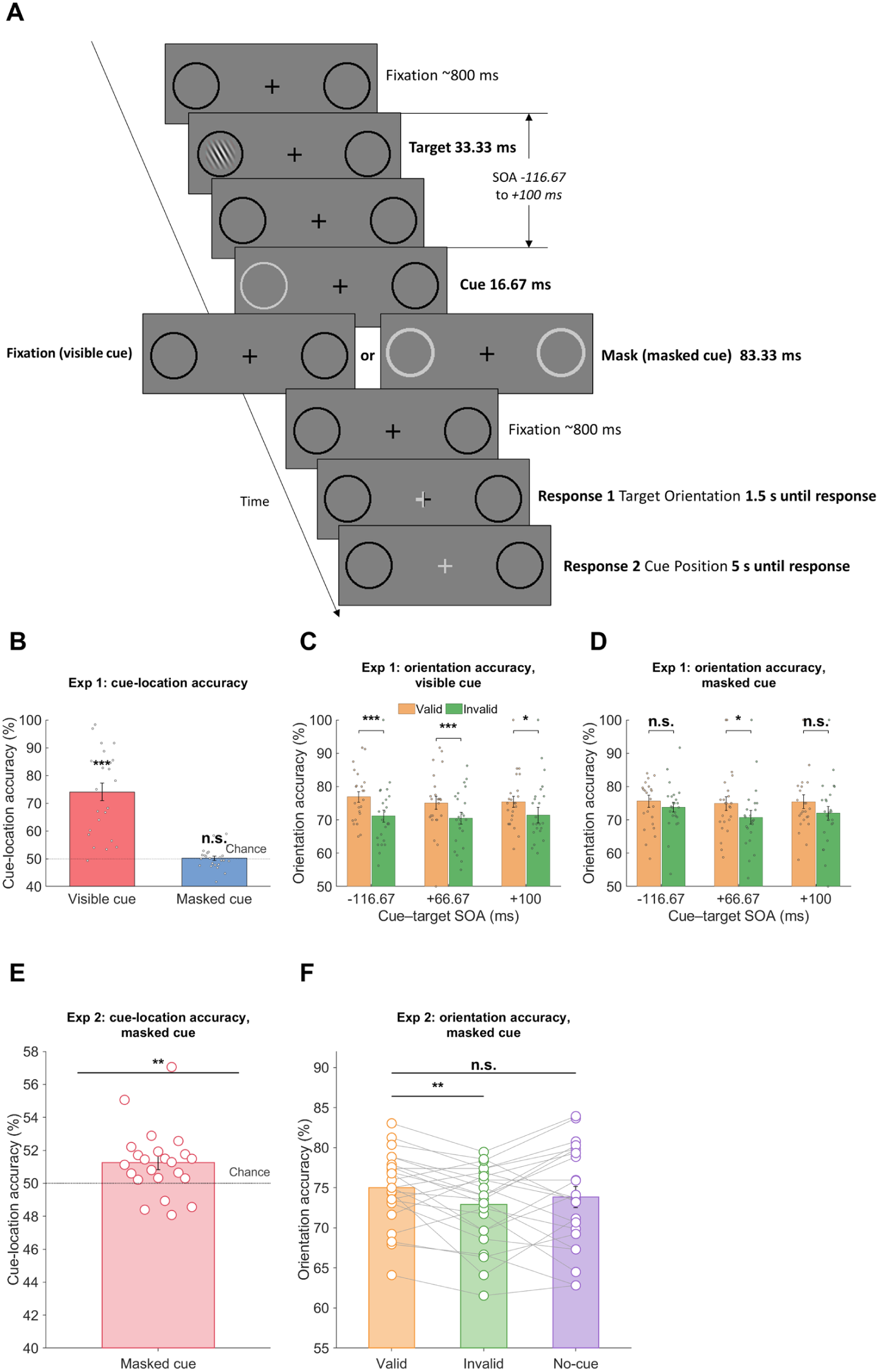
Experiment 1 and 2 behavioral results. (A) Experiment 1 trial sequence. (B-D) Experiment 1 cue-location and orientation-discrimination results. (E) Experiment 2 masked cue-location accuracy (N = 23), with the dotted line denoting chance. (F) Experiment 2 orientation accuracy for Valid, Invalid, and No-cue trials. Bars show group means, circles show individual participants, and error bars denote SEM. See also S1 Fig and S2 Fig, and S1 Table and S2 Table.

Experiment 2 (N = 23) reproduced the benefit at the +66.67 ms interval in an independent sample with masked cues (Valid 75.00% versus Invalid 72.91%, t(22) = 2.88, Holm-adjusted p = .026; No-cue 73.85%; Fig 2F), with cue-location accuracy again within ±5 percentage points of chance (51.25%; Fig 2E). The target-locked N2pc was larger for Valid than for Invalid and No-cue trials (Holm-adjusted p = .012 and .045; S13 Fig)[24, 25]; after cue-evoked activity was subtracted, this difference was no longer significant (p = .093; S2 Fig). The retrospective cueing benefit was thus present in both experiments and persisted when cue-location information was strongly reduced; Experiment 3 asked at which processing stage it operates.

### A neural measure of target evidence tracks visibility and discrimination

Having established the behavioral benefit of retrospective cueing at the critical interval, Experiment 3 asked at which processing stage this benefit arises: on the lingering sensory trace, on a report-independent target representation, or on decision and report processes (Fig 1). The three accounts make distinct predictions: a sensory-interaction account predicts an early cueing effect on target evidence, a report-contingent decisional account predicts no effect once explicit target report is removed, and a representational account predicts a late enhancement of target evidence that persists without report and predicts perception.

Experiment 3 included 20 participants who completed both Report and No-report sessions. Targets occurred above or below fixation at individually calibrated low, medium, and high contrast, and the retrospective cue followed target onset by 66.7 ms. In the Report session, participants judged target visibility (seen/unseen) and orientation; in the No-report session, they made no judgment about the targets and instead responded to sparse, unrelated fixation-cross events and occasional thought probes (Fig 3; Methods). In both sessions, all responses and task events occurred on a response screen that appeared about 970 ms after target onset, after the EEG analysis window.

**Fig 3.**
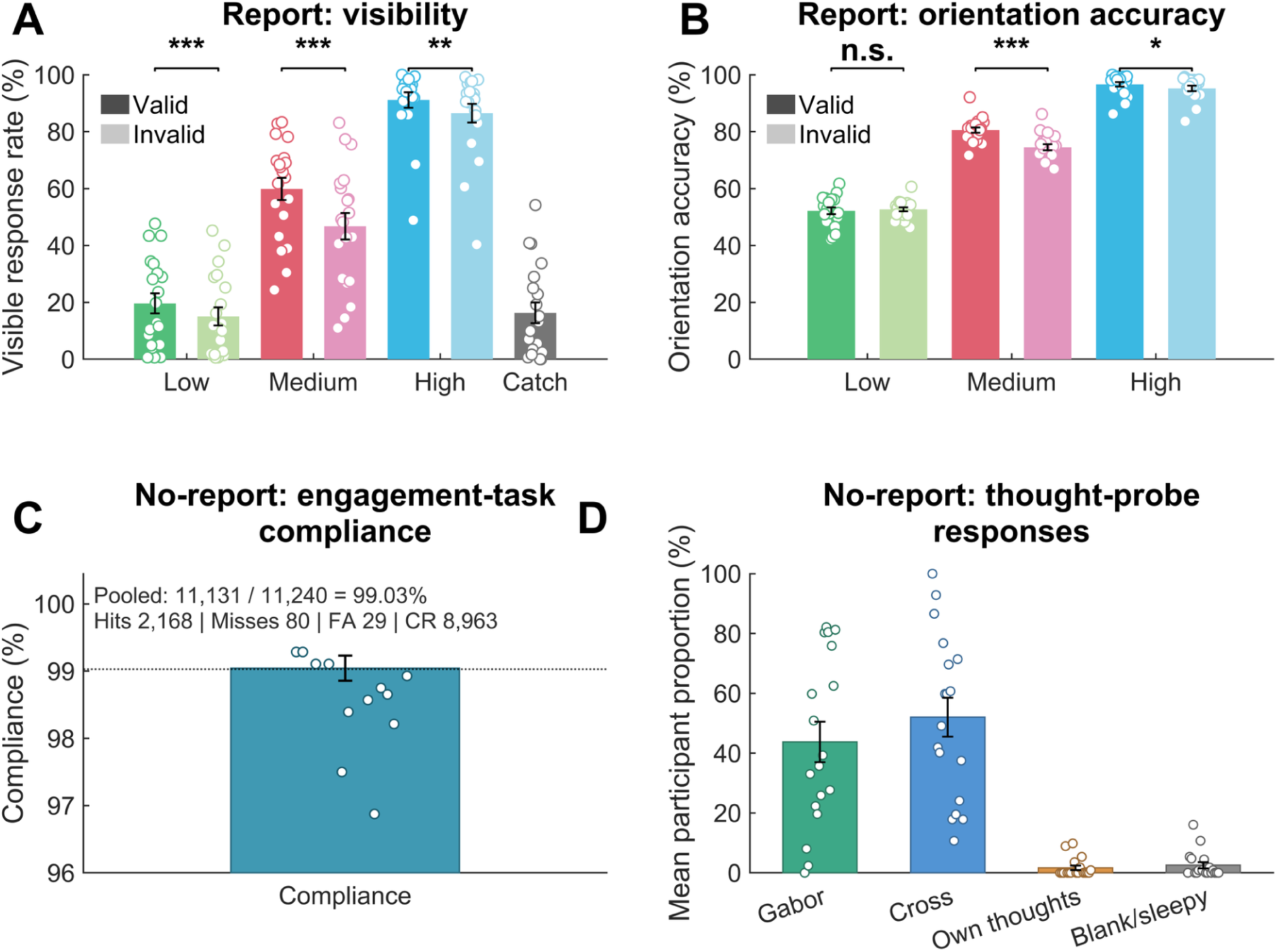
Experiment 3 behavior. A, Report binary visibility, including Catch trials. B, Report orientation accuracy. Valid and Invalid conditions are shown at Low, Medium, and High contrast; significance annotations show Valid–Invalid tests with Holm-adjusted p values (S3 Table). C, No-report engagement-task compliance; points show participant compliance, the bar and error bar show the group mean ± SEM, and the dotted line denotes pooled compliance (11,131/11,240 = 99.03%; S7 Table). D, No-report thought-probe response distribution across Gabor, cross, own thoughts, and blank/sleepy categories, summarized for N = 18 participants with at least one valid probe response; bars and error bars show group means ± SEM and points show participants (S7 Table). Of 2,248 probes, 244 were missing or invalid and are not plotted. Bars in A–B show participant means ± SEM; points show all 20 participants. See also S3 Fig and S3 Table and S7 Table.

In the Report session, orientation accuracy rose from approximately 52% at low contrast to 78% at medium and 96% at high contrast, and Seen responses rose from approximately 17% to 53% and 89%; catch trials were reported as seen on 16.6% of trials.

Valid retrospective cues increased orientation accuracy most clearly at medium contrast (80.58% valid versus 74.51% invalid); low-and high-contrast differences were smaller because performance approached floor and ceiling, respectively (Fig 3B; paired tests in S3 Table).

Visibility showed the same pattern: valid cues increased Seen responses at low, medium, and high contrast, most strongly at medium contrast (59.86% versus 46.73%). On cue-only catch trials, cue location alone did not bias Seen responses or orientation choices (both Holm-adjusted p = .888; S3 Fig C–D; S3 Table).

In the No-report session, participants remained engaged with the sparse orthogonal task (Fig 3C–D). The speeded engagement task yielded 11,131 compliant responses out of 11,240 events (99.03%; 2,168 hits, 80 misses, 29 false alarms, and 8,963 correct rejections). Thought probes yielded mean participant proportions of 43.74% Gabor, 52.03% cross, 1.69% own thoughts, and 2.55% blank/sleepy among 18 participants with at least one valid response; the Gabor was thus often reported as recent conscious content, although it was never the object of a trial-wise report.

The primary decoder was trained to distinguish High-contrast target-present from cue-only trials, with Valid and Invalid targets pooled in the target class, so that it was neutral with respect to cue validity. Its held-out signed decision value, D, measured how strongly the EEG on each trial expressed the target-present pattern (Methods). We refer to this target-present pattern as the target representation; it is not specific to target orientation.

Targets were decoded from cue-only trials at Medium and High contrast during Report and most clearly at High contrast during No-report (AUC time courses in S4 Fig A–B). In the Report session, Medium-contrast Valid-minus-Invalid D formed a corrected cluster from 352– 608 ms (cluster p = .0002), whereas at High contrast, where Report performance approached ceiling, no corrected cluster formed (Fig 4A); the No-report effect is described below (Fig 4B; all clusters in S4 Table).

**Fig 4.**
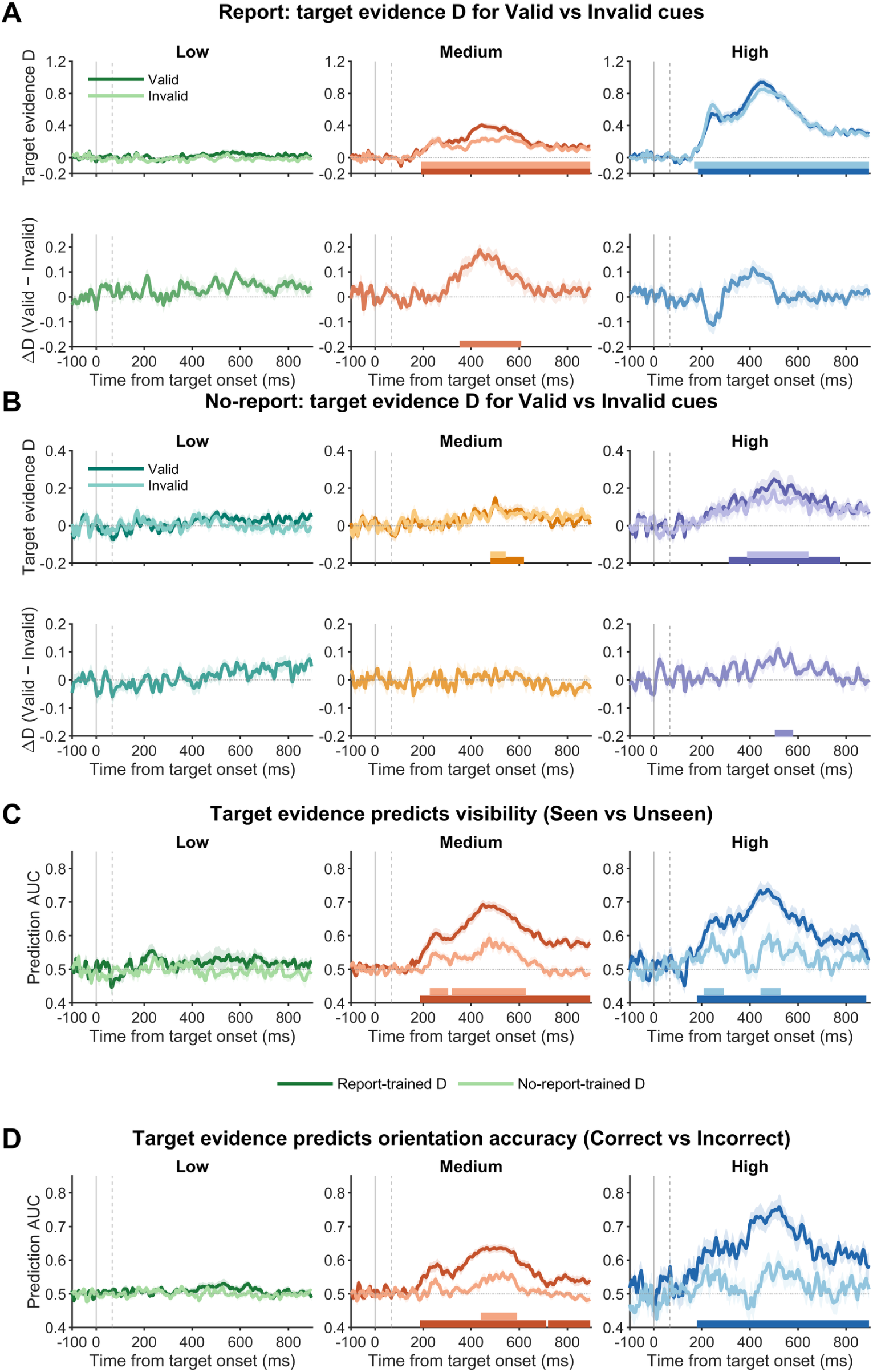
Target-related neural evidence and its relation to perceptual outcomes. A–B, held-out target evidence D from the validity-neutral Balanced decoder during Report and No-report, respectively. For each contrast, upper panels show Valid and Invalid D effects and lower panels show Valid-minus-Invalid differences; shading shows SEM, target onset is 0 ms, and cue onset is 66.7 ms. Horizontal bars denote corrected participant-level cluster-permutation intervals: in the upper panels, Valid and Invalid D-versus-zero intervals from the six-curve Low/Medium/High by Valid/Invalid correction family; in the lower panels, Valid-minus-Invalid ΔD intervals from the separate three-contrast correction family. In Report, Medium ΔD formed a corrected 352–608 ms cluster (cluster p = .0002), whereas High ΔD had no corrected cluster. In No-report, High ΔD formed a corrected 504–580 ms cluster (cluster p = .0054). C– D, time-resolved prediction of Report Seen responses (C) and orientation accuracy (D) from both Report-trained and No-report-trained D; horizontal bars denote corrected prediction intervals. Within-task AUC time courses and the No-report High 300–600 ms participant summary are shown in S4 Fig A–C. See also S4 Fig, S7 Fig, and S8 Fig, and S4 Table and S5 Table.

In mixed-effects models including late D, contrast, validity, and their interactions, the simple slope of D was positive in the High-contrast Valid reference condition: greater late D increased both the probability of a Seen response (β = 1.023, OR = 2.78, 95% CI [2.22, 3.48], p = 7.38 × 10⁻¹⁹) and orientation accuracy (β = 0.837, OR = 2.31, 95% CI [2.02, 2.64], p = 2.67 × 10⁻³⁴). D also predicted accuracy after controlling for Seen (β = 0.642, OR = 1.90, 95% CI [1.66, 2.17], p = 1.26 × 10⁻²⁰). Subject-level AUCs averaged across contrasts and the late window were above chance for Seen (mean AUC = .625, one-sided p = 2.68 × 10⁻⁸; Fig 4C) and orientation Correct responses (mean AUC = .590, one-sided p = 1.03 × 10⁻⁸, 20/20 participants; Fig 4D). Thus, D tracked trial-by-trial variation in a target representation linked to both subjective visibility and objective orientation discrimination.

### Valid cues enhance target evidence after 300 ms, even without explicit report

Under No-report, target evidence was measured relative to cue-side-matched cue-only trials and averaged equally across target locations. At High contrast, the corrected Valid-minus-Invalid cluster spanned 504–580 ms (cluster p = .0054). In the 300–600 ms late window, Valid exceeded Invalid across participants (mean ΔD = .0505, 95% CI [.0107, .0903], t(19) = 2.65, Holm-adjusted one-sided p = .0235, dz = .59; 15/20 participants positive; Fig 4B; S4 Fig C). The all-trials sensitivity decoder showed the same effect (mean ΔD = .0579, Holm p = .0217, cluster 504–580 ms, cluster p = .0038). Excluding trials that followed a thought probe left the effect unchanged (ΔD = .0595, t(19) = 2.78, two-sided p = .012; S8 Table). Thus, valid retrospective cues strengthened target evidence without explicit report. No enhancement was present early: in the 200–300 ms window, Valid and Invalid target evidence did not differ (mean ΔD = .013, t(19) = 0.86, p = .40). The late effect formed a cluster over right-hemisphere electrodes, mostly posterior and temporal (12 electrodes, cluster p = .037; S12 Fig). In Report, the validity effect was larger in the late than in the early window at Medium (late-minus-early ΔD = .118, t(19) = 5.35, Holm-adjusted p = .0002, dz = 1.20; 18/20 participants) and High contrast (.104, t(19) = 4.00, Holm-adjusted p = .004, dz = 0.89), with no early effect at Medium contrast (ΔD = .001, p = .97; S9 Fig). Searchlight decoding showed that, where early cueing effects were present, they were reversed: Invalid exceeded Valid over posterior electrodes in Report at Medium (14 electrodes, cluster p = .009) and High contrast (46 electrodes, cluster p = .002), with no early cluster under No-report (S12 Fig). An early, additive sensory interaction between cue and target would raise early target evidence; instead, the early effect was absent or reversed, and the corrected enhancement appeared only after 300 ms, consistent with attentional amplification of an already-encoded target representation.

### No-report-trained neural evidence predicts behavior in the Report session

To test whether the No-report target representation carries information about perception, we applied the No-report-trained decoder to trials of the separate Report session without retraining. No-report-derived late D predicted Seen responses (β = .368, OR = 1.44, 95% CI [1.22, 1.71], p = 2.21 × 10⁻⁵), Correct responses (β = .319, OR = 1.38, 95% CI [1.22, 1.55], p = 2.07 × 10⁻⁷), and Correct responses after controlling for Seen (β = .230, OR = 1.26, 95% CI [1.12, 1.42], p = 1.47 × 10⁻⁴). Subject-level cross-task AUCs averaged across contrasts and the late window were above chance for Seen (mean AUC = .539, one-sided p = 2.93 × 10⁻⁴, 17/20 participants; Fig 4C) and Correct (mean AUC = .521, one-sided p = 7.30 × 10⁻⁴, 17/20 participants; Fig 4D). No such prediction was present before target onset, arguing against a nonspecific association between sessions. Thus, a neural target representation learned entirely under No-report predicted both subjective visibility and objective orientation discrimination in the separate Report session.

### Report and No-report share a target representation, and Report adds further information

Cross-task target decoding was above chance at Medium and High contrast in both directions. At High contrast, late Report-to-No-report AUC averaged .553 for Valid and .534 for Invalid trials (Holm-adjusted p = .00251 and .0209), whereas No-report-to-Report AUC averaged .568 and .573, respectively (both Holm-adjusted p = .00151). Thus, target-related representations generalized across report requirements in both directions (Fig 5A–B). Full cluster statistics are reported in S4 Fig and S4 Table.

**Fig 5.**
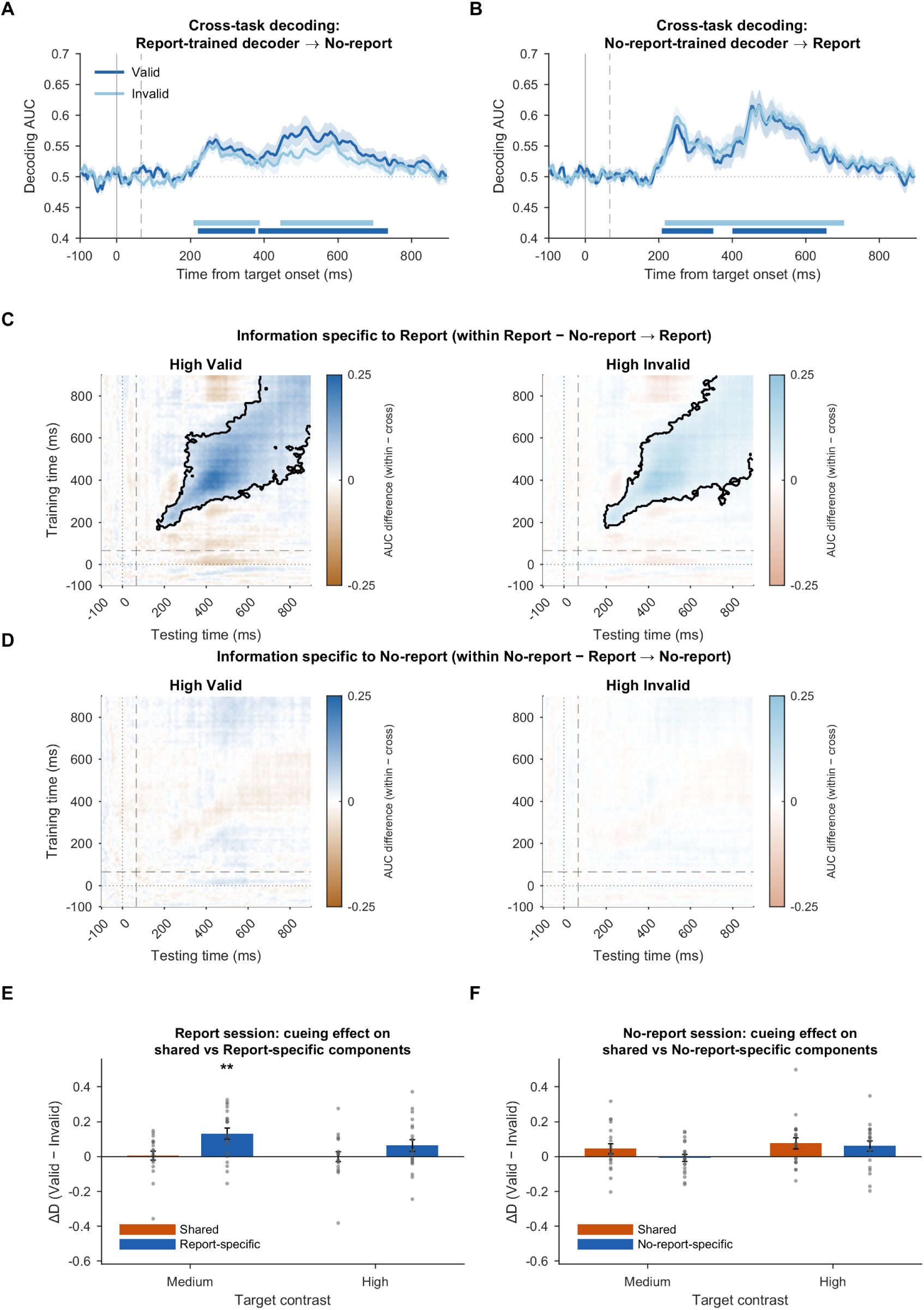
Shared and task-specific target representations across report requirements. A–B, cross-task decoding with the validity-neutral decoder. A, Report-trained decoders tested on No-report trials. B, No-report-trained decoders tested on Report trials. High-contrast Valid and Invalid curves are shown for both directions; shading shows SEM, solid and dashed vertical lines mark target and cue onset, and dark and light horizontal bars mark the corrected Valid and Invalid AUC clusters, respectively. C–D, High-contrast task-specific target information quantified as within-task minus the corresponding cross-task temporal-generalization matrix while holding the test task fixed. C, Report fixed as the test task (within Report minus No-report-to-Report), showing corrected positive specificity clusters for both Valid and Invalid trials. D, No-report fixed as the test task (within No-report minus Report-to-No-report), showing no additional No-report-specific information. E–F, cueing effects (Valid − Invalid ΔD, 300–600 ms) on the shared component (orange) and the session-specific component (blue) for Medium and High targets. E, Report session: the Report decoder split into a component along the No-report decoder axis (shared) and a component orthogonal to it (Report-specific). F, No-report session: the No-report decoder split in the same way along the Report decoder axis (shared) and orthogonal to it (No-report-specific). Bars show mean ± SEM and dots show participants (N = 20); **, Holm-adjusted p < .01 (family of four per session). Black contours denote corrected matrix clusters. Full cross-task matrices and all contrasts are provided in S4 Fig D–E; full task-specificity component matrices and participant-level specificity summaries are provided in S5 Fig. See also S4 Fig, S5 Fig, S6 Fig, S7 Fig, and S11 Fig, and S4 Table, S5 Table, and S8 Table.

We next asked whether either task contained target-related information beyond the representation shared across Report and No-report. For each fixed test task, we compared within-task temporal generalization with the corresponding cross-task generalization (Fig 5C–D; S5 Fig), following the logic of previous cross-task decoding work[16, 20, 21]. When Report was the fixed test task, within-task decoding exceeded No-report-to-Report generalization at

High contrast for both Valid and Invalid trials (late-window stability within-minus-cross differences = .0932 and .0858, respectively; both Holm-adjusted p < .00001), with corrected matrix-level specificity clusters for both conditions (cluster p = .0002). In contrast, when No-report was the fixed test task, the corresponding within-minus-cross differences were close to zero (Valid: ΔAUC = −.0077, Holm-adjusted p = .730; Invalid: ΔAUC = −.0023, Holm-adjusted p = .799). Thus, Report showed target-related information beyond what transferred from the No-report decoder, consistent with additional report-associated information, whereas the target-related representation expressed during No-report was largely captured by the Report-trained decoder.

### Report amplifies the cueing effect along report-specific dimensions

To locate the cueing effect relative to this shared representation, we split the Report decoder into a component along the No-report decoder axis (shared) and a component orthogonal to it (Report-specific; Methods). At Medium contrast, Valid exceeded Invalid on the Report-specific component (mean ΔD = .131, 95% CI [.063, .198], t(19) = 4.06, Holm-adjusted p = .003, dz = 0.91; 16/20 participants), whereas the shared component showed no validity difference (ΔD = .005, t(19) = 0.19, Holm-adjusted p = 1; Fig 5E). Time-resolved decomposition placed this effect at 356–612 ms on the Report-specific component (cluster p = .0002), with a further High-contrast interval at 396–488 ms (cluster p = .021). The effect was larger on the Report-specific than on the shared component (difference = .126, t(19) = 2.65, Holm-adjusted p = .031). The same held when the shared direction was regressed out of the features before retraining (Report-specific ΔD = .133, Holm-adjusted p = .002; difference from shared, Holm-adjusted p = .026); when the shared axis was instead defined by the activation pattern, both components carried part of the effect (Report-specific ΔD = .109; shared ΔD = .059, Holm-adjusted p = .049; S8 Table). The Report-specific component was not an unstructured residual: its activation pattern[39] was positive over central and parietal electrodes (23-electrode cluster, p = .0002) and resembled the Report-session P3b topography (mean r = .85, 20/20 participants), more closely than the shared component did (mean r = .46; difference, Holm-adjusted p = .0006; S12 Fig). It was also the stronger predictor of behavior: in joint models it predicted Seen responses (β = .58, OR = 1.78, p < 10⁻¹²) and orientation accuracy (β = .47, OR = 1.60, p < 10⁻²³), whereas the shared component made a smaller independent contribution (Seen: β = .08, p = .035; accuracy: β = .09, p = .026). When the No-report decoder was decomposed in the same way, with the Report decoder axis as the shared direction, the High-contrast cueing effect was expressed on the shared component (time-resolved cluster 452–608 ms, cluster p = .002; Fig 5F; S11 Fig).

When the Report-trained decoder was applied to both sessions, the contrast dependence of the cueing effect differed between them: the Session × Validity × Contrast interaction was significant (F(2, 26020) = 5.04, p = .0065), with the No-report effect confined to High contrast (ΔD = .058) and the Report effect peaking at Medium contrast (ΔD = .118). Across sessions and contrasts, the cueing effect followed an inverted-U function of late target decodability (quadratic b = −3.21, 95% CI [−5.22, −1.20], p = .002; peak AUC = .636, 95% CI [.613, .663]), and at matched decodability it remained larger in Report than in No-report (b = .021, p = .016; S10 Fig). Thus, the cueing effect scaled with available target evidence in both sessions and was further amplified when report was required. Session order did not change the No-report cueing effect (Report-first versus No-report-first, p = .36) or the prediction of Seen responses from No-report-trained D (D × Order, p = .37), whereas the Report cueing effect was larger in participants who completed the Report session first (Order × Session × Validity, participant-level p = .049). Follow-up analyses within each order group showed that, when Report was the first session, its cueing effect exceeded the No-report effect (Report − No-report ΔD averaged over Medium and High contrast = .095, t(9) = 3.47, p = .007) and, at Medium contrast, was carried by the Report-specific component (specific − shared = .234, t(9) = 3.78, p = .004); when Report followed the No-report session, neither difference was present (.007, p = .83; .018, p = .75). The No-report cueing effect was similar whether that session came first or second (ΔD = .038 and .026; S8 Table).

Together, retrospective cues strengthened a report-independent target representation when no report was required, whereas explicit report, when performed first, amplified the cueing effect along report-specific dimensions that carried most of the trial-wise information about visibility and discrimination.

### Complementary ERP evidence

ERP validity effects depend on report, whereas contrast sensitivity does not VAN was quantified from 240–280 ms over bilateral posterior electrodes. In the Report session, High Valid and High Invalid trials differed (mean paired difference = 0.49 µV, t(19) = 4.48, Holm-adjusted p < .001, dz = 1.00), with High Invalid trials producing the more negative response. VAN also varied from low to high contrast (Holm-adjusted p < .001; Fig 6A). Thus, during report, posterior target-related activity was sensitive to both stimulus strength and cue validity.

**Fig 6.**
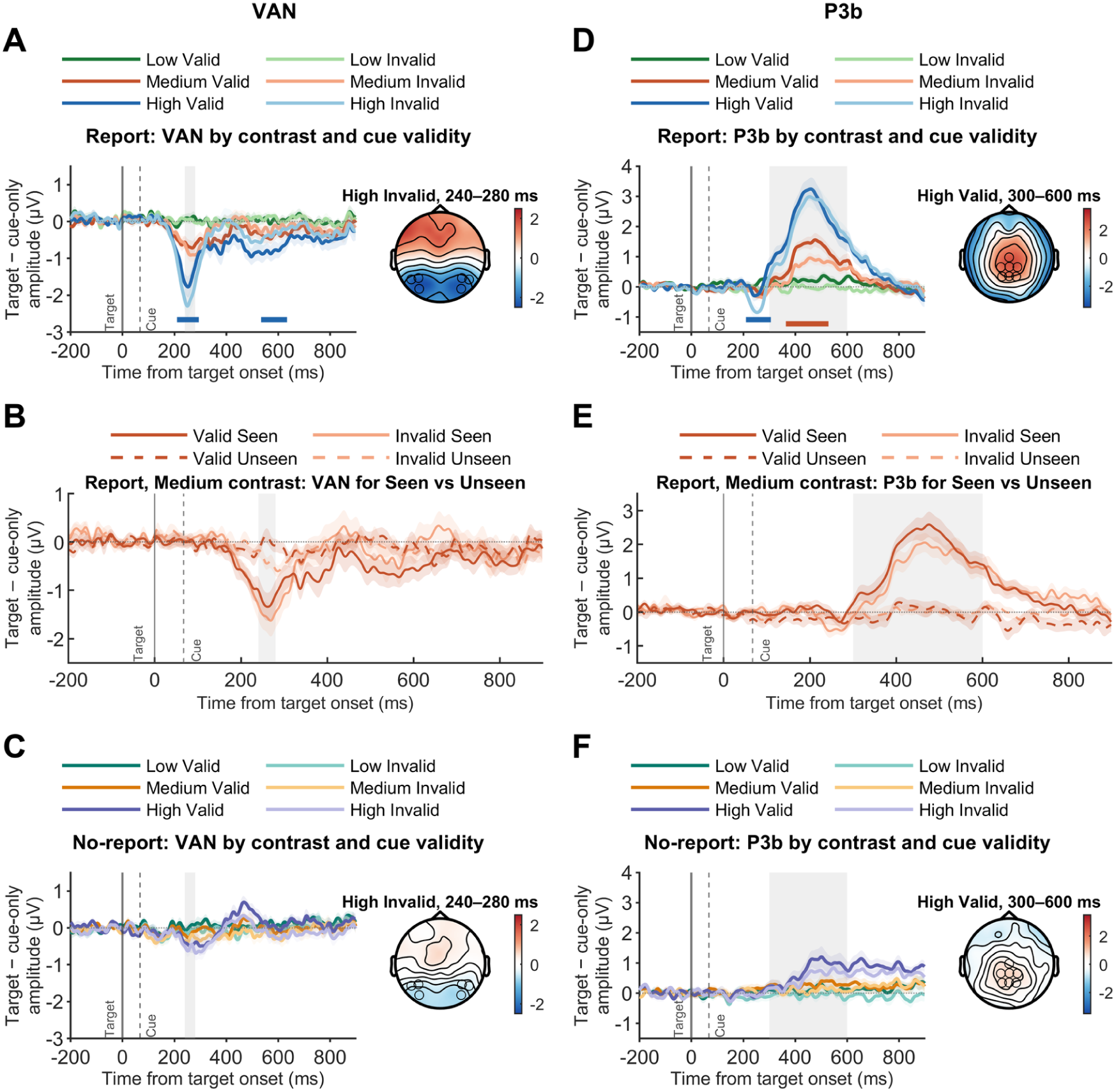
Experiment 3 cue-controlled ERP signatures: VAN and P3b. A, Report-session cue-controlled target-related VAN difference waveform (target-present minus cue-only) with the corresponding descriptive fixed-window topography. B, Medium-contrast Report-session VAN waveforms for Valid Seen, Valid Unseen, Invalid Seen, and Invalid Unseen trials. C, No-report cue-controlled VAN difference waveform with the corresponding descriptive fixed-window topography. D-F, corresponding P3b panels: D, Report-session waveform and topography; E, Medium-contrast Report-session Seen/Unseen waveforms; F, No-report waveform and topography. Waveform shading shows SEM; gray regions mark the fixed 240–280 ms VAN and 300–600 ms P3b windows; solid and dashed vertical lines mark target onset (0 ms) and cue onset (66.7 ms). Horizontal bars denote corrected time clusters. Participant-level fixed-window amplitude summaries are shown in S3 Fig E–J. See also S3 Fig.

In the No-report session, VAN remained sensitive to contrast (low-to-high mean marginal difference = -0.47 µV, t(19) = -3.55, Holm-adjusted p = .0086, dz = -0.79; Fig 6C), whereas fixed-window Valid-versus-Invalid contrasts did not survive correction.

P3b was quantified from 300–600 ms over a centro-parietal ROI after subtracting cue-side-matched cue-only activity. In the Report session, Medium Valid trials produced a larger P3b than Medium Invalid trials (mean paired difference = 0.40 µV, t(19) = 4.01, Holm-adjusted p = .0023, dz = 0.90). P3b also increased strongly from low to high contrast (mean marginal difference = 1.98 µV, Holm-adjusted p < .001; Fig 6D). Thus, retrospective cue validity selectively amplified late centro-parietal activity when participants reported the target.

In the No-report session, fixed-window Valid-versus-Invalid P3b contrasts did not survive Holm correction, whereas P3b still increased from low to high contrast (mean marginal difference = 0.57 µV, t(19) = 3.05, Holm-adjusted p = .0266, dz = 0.68; Fig 6F). Consistent with the report dependence of P3b[12, 17], the univariate validity effect was confined to Report, whereas late target-related processing persisted without report.

Together, cue-validity effects on VAN and P3b were seen with report, whereas contrast-sensitive target processing persisted without it.

### VAN and P3b amplitudes distinguish Seen from Unseen targets

To determine whether VAN and P3b tracked subjective visibility rather than physical contrast alone, we compared Seen and Unseen Medium-contrast trials. Across validity conditions, Seen targets elicited a more negative VAN (β = −1.17 µV, 95% CI [−1.73, −0.62], t(19.34) = −4.42, Holm-adjusted p < .001, dz = −1.01; Fig 6B; S3 Fig G) and a larger P3b (β = 1.42 µV, 95% CI [0.97, 1.86], t(19.52) = 6.68, Holm-adjusted p < .001, dz = 1.54; Fig 6E; S3 Fig J). Seen– Unseen differences were significant within both Valid and Invalid trials for both components (all Holm-adjusted p ≤ .0016).

Seen–Unseen differences were comparable for Valid and Invalid trials (Validity × Seen interaction: VAN Holm-adjusted p = .166; P3b Holm-adjusted p = .076). These visibility effects held in the stricter trial-count sensitivity analysis, and complementary time-resolved analyses yielded corresponding VAN and P3b clusters (Supporting Information). Thus, at matched physical contrast, both VAN and P3b distinguished consciously Seen from Unseen targets, linking VAN and P3b to subjective visibility independently of target contrast.

These ERP results complement the decoding results (Figs 4 and 5): retrospective attention strengthens perceptual target representations that generalize across report requirements, whereas explicit reporting engages additional task-associated neural dynamics.

## Discussion

Our findings show that retrospective spatial attention strengthens a report-independent target representation that predicts visibility and discrimination, and that explicit report amplifies this effect. Using masked retrospective cues (Experiments 1 and 2) and parallel Report and No-report sessions (Experiment 3), we tested three accounts of where a post-stimulus cue operates (Table 1) and found three linked effects. First, masked retrospective cues improve visual discrimination even when participants can barely locate the cue. Second, cueing enhances target evidence after the window of sensory encoding, without the early enhancement expected from an early, additive sensory interaction; without explicit target report, valid retrospective cues still strengthen target evidence, and a target representation learned entirely under No-report predicts trial-by-trial visibility and orientation discrimination in the separate Report session, arguing against a purely report-contingent decisional account. Third, the target representation is shared between Report and No-report, with decoders transferring in both directions, and when a report is required the cueing effect is amplified in additional report-specific signals, consistent with a representational account in which report adds a further stage of amplification. Together, these findings link attention directed after stimulus offset directly to visual perception.

### From retroperception to attentional selection

Sergent and colleagues established retroperception in a near-threshold Gabor paradigm: post-target cues improved orientation discrimination, and a subsequent experiment showed increased subjective visibility under explicit report[1]. Experiments 1 and 2 extend this phenomenon to an earlier retrospective interval and to backward-masked cues: valid cues improved objective orientation discrimination across cue intervals in Experiment 1 and at the critical +66.67 ms interval in Experiment 2, including when masking reduced cue-location accuracy to near chance. These experiments establish the behavioral phenomenon; the representational inference rests on Experiment 3.

### Post-stimulus attention strengthens perceptual target representations

Previous No-report studies showed that consciousness-linked activity can persist without an immediate perceptual report, but they manipulated stimulus strength or visibility rather than retrospective attention[16, 17, 20]; none tested whether a valid cue strengthens a target representation measured by a decoder that ignores cue validity.

Here, cue validity was manipulated after target onset, and the primary decoder was trained without regard to cue validity, so its evidence score measured a target-present pattern rather than a Valid–Invalid difference. During Report, trials with stronger target evidence were more often seen and correctly discriminated, and Valid cues enhanced this evidence at Medium contrast. Under No-report, Valid cues enhanced target evidence at High contrast. Crucially, a decoder trained in the No-report session predicted Seen responses and orientation accuracy in the separate Report session without retraining. Together, these results extend No-report evidence from stimulus strength to retrospective attention: even without report, the cue strengthens a target representation that predicts visibility and discrimination.

The ERP findings complement the multivariate results. In Report, both VAN and P3b distinguished Seen from Unseen Medium-contrast targets, whereas under No-report, ERP amplitudes still increased with contrast but showed no significant cue-validity effect. Thus, report requirements reshape classic ERP components[19, 29], whereas the report-independent cueing effect is detected by multivariate decoding. This fits the finding that the P3b and temporally stable late decoding depend on report[17]: the No-report cueing effect was a late increase in target evidence that did not generalize stably over time (S8 Fig).

### Shared representations and report-associated processing

Cross-task decoding revealed a target representation shared between sessions, plus additional report-related information. At high contrast, decoders trained in one session detected targets in the other, in both directions. Task-specificity analysis showed that Report trials contained target information that the No-report-trained decoder did not capture (Fig 5C–D; S5 Fig). During Report, the cueing effect was carried mainly by this extra, report-specific information (Fig 5E), whose scalp pattern resembled the P3b (S12 Fig). Sergent and colleagues found a similar asymmetry between active and passive listening[16]: a decoder trained on active sessions captured most of the late activity in passive sessions, but active sessions contained extra information that a passive-trained decoder missed. Sergent and colleagues proposed that late activity without a task reflects conscious access itself, in which a representation is maintained and shared across a broad brain network (a “global playground”), and that an explicit task adds a decision-related P300 component, turning this playground into a global workspace[16]. Our results show a similar division: a shared representation present without report, and P3b-like report-specific signals added when a report is required. The parallel concerns this split between shared and task-added signals; the bifurcation dynamics that Sergent and colleagues used to identify conscious access were not tested here. Our results extend this idea to attention: the retrospective cue strengthens the shared representation, and explicit report amplifies its effect through report-specific signals.

The cueing effect also peaked at different contrasts: at Medium contrast in Report, but at High contrast in No-report. One explanation is that a task-irrelevant target is processed more weakly[30]: in No-report, High-contrast targets were decoded about as well as Medium-contrast targets in Report (mean AUC = .547 versus .580; Report High, .716). Across sessions, the cueing effect was largest for targets of intermediate decodability (an inverted-U relation; S10 Fig). At matched decodability, the effect remained larger in Report, consistent with amplification by report. This amplification depended on session order: it was clear when the Report session came first and absent when it followed the No-report session, whereas the No-report effect did not depend on order. One possibility is that the additional benefit under report declines once the stimuli have become familiar, whereas the report-independent effect is stable.

### Theoretical implications and scope

At the theoretical level, the findings support the idea that attention can still strengthen a sensory representation after the stimulus has gone, and that this representation is linked to what participants report seeing[1, 2, 40]. This fits workspace theories in which attention controls which content gains access[8, 9, 31, 32], and the global-playground proposal, in which a representation is maintained without a task and is joined by decision processes when a report is required[16]. Our results identify the stage at which the retrospective cue operates; they do not decide between broad theories of consciousness. The larger P3b and the extra report-specific information during Report further indicate that reporting adds late, task-related processing on top of the shared representation, and that this processing amplifies the effect of retrospective attention when the Report session comes first.

Two design features support this interpretation. First, masking reduced cue-location accuracy to near chance, and Experiment 2 reproduced the behavioral benefit in a new sample. Second, the No-report session followed the design of Sergent and colleagues[16]: task events and thought probes appeared on the response screen after the analysis window, and were unrelated to target presence and cue validity; together with cross-task decoding, this minimized the contribution of explicit target report, a general challenge for no-report paradigms[33]. Overall, retrospective attention strengthens a report-independent visual target representation that predicts visibility and discrimination, with no detectable modulation by session order; explicit report adds a further amplification when the Report session comes first.

## Materials and Methods

### Experimental Model and Study Participant Details

A total of 29 human subjects (9 male; age range, 18–28 years) were recruited for Experiment 1. The primary analysis retained 24 participants; five participants (4 male, 1 female) were excluded because their overall behavioral accuracy in the formal experiment fell below 65% or exceeded 85%, that is, outside the range around the 75% staircase target used for inclusion. The retained cohort therefore comprised 24 participants (5 male, 19 female). Twenty-three human subjects (9 male; age range, 18–29 years) participated in Experiment 2, and all 23 were retained under the unified criterion requiring clean EEG for at least 70% of valid behavior trials (minimum retained = 74.11%). Twenty human subjects (9 male; age range, 18–29 years) participated in Experiment 3, and no subject was excluded. Sample sizes were based on previous studies of retrospective attention[1, 2] and an a priori power calculation using G*Power (RRID:SCR_013726)[34], which indicated that samples of 24, 23, and 20 participants would provide 0.80 power to detect a medium-sized within-subject effect (f = 0.35). Because the main inferences rely on mixed-effects models and participant-level permutation tests rather than on this ANOVA, we report effect sizes, confidence intervals, and the number of participants showing each effect alongside the tests. Participants were naive to the purpose of the study, except for one participant in Experiment 3 who was an author; excluding this participant left the No-report cueing effect, the cross-task prediction of Report behavior, and the decoder decomposition unchanged in direction and significance (all N = 19 results in S8 Table). All participants were right-handed, reported normal or corrected-to-normal vision, and had no known neurological or visual disorders. They provided written informed consent, and all procedures and protocols were approved by the Human Research Ethics Committee for Non-Clinical Faculties of the School of Psychology at South China Normal University (approval reference number: SCNU-PSY-2023-043). Data were collected from 6 May 2023 to 7 December 2023 for Experiment 1 and from 28 October 2023 to 7 December 2023 for Experiment 2. Data for Experiment 3 were collected from 15 May 2024 to 1 October 2024.

## METHOD DETAILS

### Stimuli

In both Experiment 1 and Experiment 2, the stimuli consisted of two horizontally arranged black circles (1.3 cd/m², 3° visual angle in diameter) on the screen, along with a central fixation cross (1.3 cd/m², 0.5° visual angle) for fixation. These circles were positioned 5° to the left and right of the fixation point. The experimental procedure was divided into a threshold-testing phase and a formal experiment phase. Cues were briefly presented (16.67 ms) as a dimming (20 cd/m²) of one of the two black circles. In the Masked cue conditions, this cue was immediately followed by an 83.33-ms bilateral dimming of both circles (with brightness adjusted individually) that served as a backward mask. In the Visible Cue conditions, no subsequent bilateral masking dimming was presented; an 83.33-ms fixation-only interval was shown instead. The target stimulus was a Gabor patch (2° visual angle in diameter) tilted either 30° or 150° from the vertical. In Experiment 3, the stimuli were similar to those in Experiment 1 and Experiment 2, with the following adjustments: the diameter of the black circles was increased to a 4° visual angle, and the diameter of the Gabor patch was increased to a 3° visual angle. Additionally, the two black circles were positioned 5° above and below the fixation point rather than to the sides.

### Behavioral experiments overview

The experiment was programmed in MATLAB R2016b (RRID:SCR_001622) using Psychtoolbox (RRID:SCR_002881) [35]. In all experiments, trial conditions were presented in random order within each block. All stimuli were presented on a 19-inch LED computer screen with a resolution of 1024×768 and a refresh rate of 60 Hz. The background was gray (28 cd/m²). The viewing distance was 57 cm, and participants’ head positions were stabilized using a chin rest.

### Threshold procedures

In Experiment 1, participants first performed a 2AFC cue-location check (left vs. right). In the visible cue condition (unmasked), accuracy was maintained above 95%, whereas in the masked cue condition (masked by an 83.33-ms bilateral dimming), backward masking strongly reduced cue-location discrimination. Target orientation discrimination thresholds were then measured without cues using a 2AFC staircase targeting 75% accuracy, completed separately for the unmasked and masked sessions (two staircases of 80 trials each; order counterbalanced across participants). In Experiment 2, only the masked cue condition was presented; participants completed cue-location checks used to quantify cue-location discrimination under backward masking and two staircase sessions (80 trials each) to determine the Gabor contrast yielding 75% orientation discrimination accuracy without attentional manipulation. In Experiment 3, retrospective cues were always unmasked (visible); participants completed a 2AFC cue-location task (above vs. below; 32 trials per run), repeated until accuracy reached at least 95%. Target contrast was then calibrated with an adaptive 2AFC staircase (80 trials per run) targeting 70%–80% accuracy. The staircase-derived Medium contrast was then used to generate the Low and High contrast levels by the predefined symmetric transformation described below, broadly spanning low (∼50% accuracy / near detection threshold), medium (∼75% accuracy), and high (∼100% accuracy) performance levels without requiring separate staircases. As described below, contrast parameters were stored separately for Report and No-report sessions.

### Experiment 1

For Experiment 1, the primary behavioral inference used all retained condition-coded trials in a binomial-logit GLMM: Correct ∼ Validity * SOA * CueVisibility + (1 + Validity | Subject), with effect coding and Laplace approximation. The random Validity slope was evaluated with full covariance, then diagonal covariance, and finally a random-intercept-only tier based on convergence and singularity diagnostics rather than p values. The four Type-III terms were Holm-corrected. The participant-level ANOVA and the masked-cue awareness analyses are reported as comparison, equivalence, and robustness analyses.

As shown in Fig 2A, each trial began with the appearance of a fixation cross at the center. After a fixation interval of approximately 800 ms, a target—a Gabor patch with a random orientation (30° or 150° away from the vertical)—was displayed for 33.33 ms within one of the circles located to the left or right of the fixation point. The target contrast was individually calibrated for each participant using a staircase method. A 16.67 ms cue, consisting of a brief dimming of one of the circles, could appear either before or after the target. This cue could be either valid (on the same side as the target) or invalid (on the opposite side). Following the cue, there was an 83.33 ms mask or fixation period for the masked and unmasked conditions.

A response screen was presented approximately 800 ms after the offset of the mask or fixation, with a response cue indicating the target’s side (marked by thickening and dimming on one side of the cross). Participants were then asked to report the target’s orientation by choosing between two options (Left/Right). Subsequently, a second response screen appeared, displaying a dimming cross at the fixation point, where participants reported the position of the cue in a two-alternative forced choice (2AFC) task.

### Experiment 2

Each participant completed two staircase sessions (80 trials each) for the masked cue condition to determine the target’s contrast at which they could discriminate the target’s orientation with 75% accuracy, without any attentional manipulation. This contrast value was then fixed for the remainder of the experiment. Participants subsequently performed four experimental blocks, each consisting of 153 trials, comprising 204 target-absent catch trials (51 trials per block, 33.33% of all trials) and 408 target-present trials, yielding exactly 136 trials for each of the three cueing conditions (Valid, Invalid, and No-cue).

The stimulus presentation for the EEG experiment was consistent with that of Experiment 1; in recordings with complete trigger logs (15 of 23), the response screen followed target onset by 1,052–1,079 ms. The procedure in Experiment 2 was fundamentally similar to Experiment 1, with the following modifications: the two circles were placed below the fixation point (shifted downward by 2.5° of a visual angle compared to Experiment 1); on 33.33% of trials, the Gabor target was omitted, forming target-absent catch trials (cue–mask sequences in the cued conditions and mask-only sequences in the No-cue condition); the stimulus onset asynchrony (SOA) between the target and the cue was fixed at 66.67 ms; a No-cue condition was added; and only the masked cue condition was retained.

### Experiment 3

Each participant completed an adaptive 2AFC target-contrast staircase (80 trials per run) targeting 70%–80% orientation accuracy without attentional manipulation. The resulting threshold contrast served as the Medium contrast level (c_Medium), from which Low and High contrast levels were defined symmetrically for each participant by subtracting and adding an individualized contrast offset (c_Low = c_Medium − Δc, c_High = c_Medium + Δc, with mean Δc ≈ 0.011). The saved Low, Medium, and High contrast values used for stimulus presentation were fixed within the corresponding formal session. Nominally, participants performed four experimental blocks, each consisting of 196 trials, comprising 112 cue-only catch trials (28 trials per block, 14.29% of all trials; 56 upper and 56 lower) and 672 target-present trials, yielding exactly 112 trials for each of the six cueing-by-contrast conditions (Low-Valid, Low-Invalid, Medium-Valid, Medium-Invalid, High-Valid, and High-Invalid).

Trial timing followed Experiment 1 (fixation interval before target onset: median 804 ms). Two circles were positioned above and below fixation. On target-present trials, the Gabor patch and retro-cue independently appeared in one of the two circles. Stimulus intensity was determined for each participant from the calibrated contrast levels: Medium contrast targeting approximately 75% orientation discrimination, with Low contrast (∼50% accuracy) and High contrast (near-100% accuracy) derived symmetrically around the Medium threshold. Cue-only catch trials contained no Gabor target; instead, an upper or lower retro-cue was presented at the same locations, with no Valid/Invalid status. The saved normalized Gabor contrast parameter was 0.00645 ± 0.00379 for Low, 0.01818 ± 0.00649 for Medium, and 0.03058 ± 0.00963 for High in No-report (mean ± SD; N = 20); corresponding Report-session values were 0.00605 ± 0.00366, 0.01762 ± 0.00556, and 0.03068 ± 0.00941. Report and No-report values were kept separate because they were not identical for all participants. Applying a luminance-meter-derived RGB gray-level-to-luminance calibration of the experimental display to the nominal dark and bright RGB endpoints implied by each saved parameter yielded luminance-domain Michelson contrasts of 0.03795 ± 0.02479 for Low, 0.10516 ± 0.03762 for Medium, and 0.17824 ± 0.05620 for High in No-report (mean ± SD; N = 20); corresponding Report-session values were 0.03666 ± 0.02277, 0.09938 ± 0.03535, and 0.17597 ± 0.05545. As a descriptive check of the spacing of the three calibrated stimulus levels, the group-mean luminance-calibrated Michelson contrasts increased approximately linearly from Low to Medium to High (R^2 = .999 for No-report and .997 for Report; S6 Table).

Each participant completed the Report and No-report sessions on the same day, in counterbalanced order (10 participants started with each session). In the Report condition, participants jointly identified target orientation (Left or Right) and subjective visibility (Seen or Unseen), yielding four response options. On cue-only catch trials, participants used the same four-option response to provide an orientation choice and Seen/Unseen judgment despite no Gabor target being presented.

Additionally, a No-report condition with sparse orthogonal engagement tasks was introduced. Task events were assigned trial-by-trial independently of the stimulus sequence via a separate random permutation, ensuring strict orthogonality to target presence, contrast, and retro-cue validity. Five sevenths of trials (∼71.4%) contained engagement-response events, with a Go to no-key-response ratio of 1:4:

1. Engagement and Speed-Response Events (5/7 of all trials, ∼71.4%): On 1/7 of all trials (∼14.3%), a green half-cross served as a speeded Go signal requiring a space-bar press within a 1-s response window. The remaining 4/7 of trials (∼57.1%) required no keypress: these comprised red No-go half-crosses on 2/7 of trials (∼28.6%; withhold response) and gray half-cross response cues pointing toward the target Gabor location on 2/7 of trials (∼28.6%; identical to the response cue in Report sessions, presented for 1 s but requiring no keypress).
2. Mind-Wandering Thought Probe (1/7 of all trials, ∼14.3%): Participants answered the intermittent self-paced question “What did you just see / what were you thinking?” by choosing one of four options: “the Gabor,” “the cross,” “my thoughts,” or “blank / I feel sleepy” (followed by a 1.5-s fixation).
3. Simple Continuation Prompt (1/7 of all trials, ∼14.3%): Participants pressed the space bar to continue when prompted (followed by a 1.5-s fixation).

This schedule yielded 784 trials per participant. In the No-report session, 18 participants contributed 784 trials, one contributed 728 trials, and one contributed 896 trials, yielding 15,736 trials in total (11,240 engagement events, 2,248 thought probes, and 2,248 continuation prompts; S7 Table). Thus, “No-report” refers here specifically to the absence of explicit perceptual reports about the Gabor targets, not to the absence of thought about them, which the probes were designed to sample; engagement tasks and thought probes, which maintained engagement and sampled ongoing conscious content, were scheduled independently of target presence, contrast, and retro-cue validity. In both sessions, all task events and responses occurred on the response screen, which appeared 970–1036 ms after target onset (median 972 ms), after the end of the EEG epoch.

### Cue-only catch-trial response-bias analysis

To test whether cue location itself induced a spatial report bias, we analyzed only Report-session cue-only catch trials (upper- and lower-cue trials) with valid four-option responses. Because no Gabor target was presented, catch trials had neither target location nor Valid/Invalid status. For each participant, upper- and lower-cue Seen proportions were compared with a two-sided paired t test. Orientation response was coded as left-tilted for bins 1/2 and right-tilted for bins 3/4 and tested at trial level with the binomial-logit mixed model RightChoice ∼ CueUpper + (1 + CueUpper | participant), fitted by Laplace approximation. The two planned cue-location tests were Holm-corrected.

### EEG data acquisition

EEG was recorded with a 64-electrode actiCAP arrangement and a Brain Products amplifier. FCz served as the online reference, AFz as ground, and an infraorbital IO channel monitored eye movements and blinks. Impedances were kept below 10 kΩ. Continuous data were acquired at 1000 Hz without online filtering.

### Data preprocessing

Offline preprocessing was performed in BrainVision Analyzer 2.2 (Brain Products; RRID:SCR_002356). Noisy channels were identified and interpolated when necessary; a 40-Hz low-pass filter and whole-scalp average reference were applied, and ocular artifacts were corrected with ICA. The ICA-corrected clean data were exported at 500 Hz with 65 recorded labels; MATLAB removed the IO channel, leaving 64 scalp channels, and applied no additional filtering, rereferencing, or ICA. Experiment 2 target-onset-aligned epochs covered -200 to 700 ms and used a -100 to 0 ms baseline. The Experiment 2 cohort gate required retained clean EEG trials to be at least 70% of valid behavior trials, complete coverage of all nine trigger-defined conditions, at least 20 trials in every N2pc condition-by-target-side cell, and at least 20 trials in every MVPA class. Experiment 3 target-onset-aligned epochs covered −200 to 900 ms and likewise used a −100 to 0 ms prestimulus baseline.

### Cue-controlled ERP difference waves

For Experiment 3, trials were mapped to canonical conditions using subject-specific trigger tables, including a predefined No-report trigger mapping for one participant. Target-present ERPs were matched to cue-only ERPs by cue location. Upper-and lower-location target-minus-cue-only difference waves were computed first and then averaged with equal weight, preventing unequal location counts from determining the result. These cue-controlled difference waves were used for the Experiment 3 VAN and P3b analyses: because the cue followed target onset by only 66.7 ms, cue-evoked responses overlap the VAN and P3b intervals, and the matched subtraction reduces this contribution.

Experiment 2 was treated differently. Its primary N2pc analysis used conventional target-locked contralateral-minus-ipsilateral difference waves without cue subtraction. A target-minus-cue-only cue-controlled N2pc analysis was included as a supplementary sensitivity analysis to assess the extent to which cue-evoked lateralized activity contributed to the primary effect.

The zero point denotes target onset; the retrospective cue occurred at +66.7 ms. Subject-level amplitudes and time courses were retained for all statistical analyses.

### ERP analysis

Component windows and regions of interest were fixed on the basis of previous studies[24–27] and applied unchanged to the corresponding analyses in Experiments 2 and 3: Experiment 2 N2pc, 240–280 ms at O1/O2, PO3/PO4, and PO7/PO8; Experiment 3 VAN, 240–280 ms at P5/P6/P7/P8/PO7/PO8; Experiment 3 P3b, 300–600 ms at Pz/P1/P2/CPz/CP1/CP2/Cz.

For the Experiment 2 N2pc, contralateral and ipsilateral ROI waveforms were averaged within left-and right-target trials and then combined with equal side weighting. The primary condition test was a one-factor repeated-measures ANOVA on each participant’s Valid, Invalid, and No-cue difference-wave means. Cue-controlled target-related N2pc served as a sensitivity analysis. A frontotemporal ocular proxy, mean(Fp2,F8,FT8) minus mean(Fp1,F7,FT7), was summarized over 0–300 ms and entered as a covariate in a participant-level N2pc model; no channel was treated as a dedicated ocular electrode.

For each Experiment 3 component, fixed-window paired contrasts were corrected with Holm’s method. ROI Valid-minus-Invalid time courses were assessed from 0–700 ms with 5,000 subject-level sign-flip permutations and max-cluster correction across the three contrasts. Component topographies are descriptive and use the condition with the largest absolute ROI amplitude within the fixed window; Report and No-report maps share symmetric zero-centered scales.

### Seen/Unseen ERP analysis

Binary visibility was defined from the four response bins (1 and 4 = Seen; 2 and 3 = Unseen). Cue-only trials retained their recorded Seen/Unseen responses for the behavioral response-bias analysis, but they were not assigned to target Seen/Unseen ERP cells; in ERP analyses they served only as participant- and cue-side-matched baselines. For each target trial, ROI amplitudes and waveforms were calculated before and after subtraction of the participant- and cue-side-specific Report cue-only mean. The confirmatory analysis was restricted to Medium contrast. Trial-level linear mixed-effects models tested residual amplitude as Seen × Validity + CueSide + (1 + Seen | participant), using effect coding and Satterthwaite degrees of freedom. Random effects were reduced to independent slopes and then random intercepts only if the richer model failed or was degenerate.

All 20 participants were retained in the primary trial-level model. A strict sensitivity analysis retained 17 participants for whom all four Medium Validity × Seen cells contained at least 15 trials and both cue locations. A three-contrast model was secondary because strict complete-cell samples were only 7 at Low and 2 at High contrast. Medium Seen-minus-Unseen ROI time courses were tested exploratorily from 0–700 ms in the strict sample using 5,000 two-sided subject-level sign flips with joint max-cluster correction across Valid and Invalid panels within each component. Fixed-window tests were the primary analyses, and Holm correction was applied within each family.

### Multivariate pattern analysis

All multivariate analyses used MVPA-Light[36]. ERP data remained at 500 Hz; MVPA inputs were anti-alias downsampled to 250 Hz, yielding a 4 ms time step. Report and No-report sessions were modeled independently. The primary validity-neutral Balanced decoder trained a shrinkage LDA to separate high-contrast target-present trials from task-matched cue-only trials, with High Valid and High Invalid trials jointly defining the target class, so that the decoder was trained blind to cue validity. Within each training fold and cue side, High Valid, High Invalid, and cue-only trial counts were balanced (Valid targets = Invalid targets; target-present total = cue-only total). Nine-fold cross-validation was repeated 25 times.

Class undersampling, feature standardization, and decision-value orientation were estimated from training data only. Undersampling was applied only within training folds; held-out test folds retained all trials without downsampling. Each anchor trial received one held-out value per repeat, and lower-contrast and High-Invalid trials were projected only by models that had not used those trials for training. Decision values were oriented target-positive using the training-data sign and scaled by the training-score standard deviation; no condition-wise z-scoring or probability transformation was used.

AUC quantified target-versus-cue-only discrimination. Continuous D was summarized as target evidence relative to cue-side-matched cue-only evidence. Report mixed models tested binary Seen/Unseen and orientation accuracy as functions of late D (300–600 ms), contrast, validity, and their interactions, with subject random effects; Correct was also tested with Seen included as a covariate. For the No-report-to-Report analysis, the No-report-trained models were projected onto independent Report trials and the same mixed models were fitted without retraining. Subject-level behavioral AUCs were computed at each time point within each contrast, averaged over the 300–600 ms window and across contrasts, and tested one-sided against .5 with participants as the unit. When a participant had no trials of one outcome class in a given contrast, that contrast was omitted from the participant’s average (Seen: 8 participant × contrast cells; Correct: 7). Report and No-report D contrasts and all window summaries also used the participant as the statistical unit. The all-trials decoder (equal-midpoint sampling sensitivity) retained all eligible training trials and served as a sensitivity analysis; the Balanced decoder was primary.

Temporal generalization trained a classifier at each 4 ms time point and tested it at every time point. The late-window stability index averaged off-diagonal AUC from 300–600 ms while excluding |train-test| <= 50 ms. Guard bands of 25 and 75 ms served as descriptive robustness checks. One- and two-dimensional cluster tests used 5,000 subject-level sign flips with max-cluster familywise correction; means were not extracted from significant clusters for a second inferential test. Fixed-window tests of late-window stability were Holm-corrected within each family.

Cross-task decoding used the same validity-neutral source-task training definition. A classifier trained in Report was tested on independent No-report target and cue-only trials, and vice versa. Test-task data never entered source-model training. Twelve AUC time courses were tested one-sided against .5 in one max-cluster family, and six Valid-minus-Invalid time courses were tested in a separate family. Early (200–300 ms) and late (300–600 ms) subject-level summaries used two-sided Valid–Invalid tests with Holm correction. Full component cross-temporal matrices are provided in the Supporting Information. To quantify task-specific information while holding the test task fixed, we subtracted the corresponding cross-task temporal-generalization matrix from the within-task matrix separately for Valid and Invalid trials. Positive within-minus-cross values therefore index target-related information expressed more strongly when training and testing occurred within the same task than could be transferred from the other task. High-contrast specificity maps are foregrounded in Fig 5C–D, with the full component matrices and participant-level late-window stability summaries provided in S5 Fig. For the High-contrast specificity maps, Report-Valid, Report-Invalid, No-report-Valid, and No-report-Invalid formed a single four-panel two-sided 2D cluster-permutation family (5,000 permutations; max-sum cluster statistic with minimal adjacency); positive and negative clusters were each tested at α/2 within the shared max-cluster family. Participant-level specificity late-window stability was the mean within-minus-cross AUC over the 300–600-ms train-by-test region excluding |train−test| ≤ 50 ms; the four High-contrast task-by-validity values were tested two-sided against zero and Holm-corrected together.

Late emergence of the cueing effect. For each participant, task, contrast, and validity, held-out D was averaged over an early (200–300 ms) and a late (300–600 ms) window after subtraction of the cue-position-matched cue-only mean. The window × validity interaction was tested as the paired difference between late and early ΔD (Valid − Invalid) for each task × contrast cell (two-sided; Holm correction over six tests).

Shared and Report-specific decoder components. A No-report decoder (shrinkage LDA; Balanced High-target versus cue-only training on 300–600 ms mean features, averaged over 25 balanced draws) defined a shared axis û. In each of the Report session’s 9 × 25 cross-validation folds, the Report decoder weight vector w_R was trained in the fold’s z-scored feature space and split into w_S = (w_R·û)û and w_P = w_R − w_S. Held-out trials were projected onto û and onto the unit vector of w_P, oriented target-positive and scaled using training data only, and cue-position-matched cue-only means were subtracted before averaging across repeats. The roles of the two sessions were exchanged for the No-report decomposition, so the shared projections in the two sessions are defined relative to different reference axes. Valid − Invalid ΔD was tested per component and contrast (two-sided; Holm family of four per session), and the two components were compared directly with paired tests of ΔD_specific − ΔD_shared (Holm family of two contrasts per session and definition). Time-resolved decompositions used the 4-ms decoders of the main analysis with max-cluster correction (5,000 permutations). Sensitivity analyses defined the shared axis by the activation pattern of the No-report decoder[39] or regressed the shared direction out of the features before retraining. Behavior was modeled with binomial GLMMs as above, entering both component scores jointly.

Target evidence and common-axis analysis. Late target decodability was the AUC of late D for targets versus cue-position-matched cue-only trials, averaged over Valid and Invalid targets. The cueing effect was modeled as ΔD ∼ Evidence + Evidence² + (1 | Participant) over 120 participant × cell values (evidence centered on the grand mean); the peak’s 95% CI was obtained from 5,000 participant bootstrap resamples, and Task was added as a sensitivity term. For the common-axis test, the Report-trained decoder was applied to both sessions, and trial-level late D was modeled as Session × Validity × Contrast with participant random intercepts and Validity slopes.

Searchlight decoding and activation patterns. For each electrode, the electrode and its neighbours (44-mm distance criterion; template electrode positions) served as features for the validity-neutral Balanced decoder (High targets versus cue-only; 200–300 or 300–600 ms means; cross-validation folds and balanced training rows as in the main analysis), and held-out decision values gave ΔD maps. For the decoder components, activation patterns A = Cov(X_train)w / (wᵀCov(X_train)w)[39] were computed in channel space in each training fold, averaged across folds and repeats, and scaled to unit norm per participant. Maps were tested across electrodes with cluster-based sign-flip permutations (5,000; sum of t; max-cluster), and spatial similarity with the Report-session P3b topography (300–600 ms, Medium contrast) was assessed with Fisher-transformed Pearson correlations.

### Use of artificial intelligence tools and technologies

During the preparation of this work, the authors used OpenAI ChatGPT, OpenAI Codex, Google Gemini, and Anthropic Claude for language refinement, manuscript organization, consistency checking, code inspection and review, debugging, documentation, verification of selected implementation details, and adjustments to graphical layout and labeling. As appropriate, the authors reviewed, edited, and verified AI-assisted outputs and retain responsibility for the manuscript, analyses, figures, and conclusions. AI tools did not generate or alter underlying research data and did not independently determine or alter the reported statistical results.

## Acknowledgments

We acknowledge the subjects for their contribution to this study. This work was supported by the National Science and Technology Innovation 2030 Major Program (2026ZD0222100), the National Outstanding Youth Science Fund Project of the National Natural Science Foundation of China (32022032), and the National Natural Science Foundation of China General Program (32271099).

## Author Contributions

X.Z. and Y.L. designed research; Y.L., X.W., M.M., L.J. and Y.C. performed research; Y.L. and X.W. analyzed data; and Y.L. wrote the paper.

## Data Availability

All processed data, figure-level source values, and author-generated analysis and plotting scripts required to reproduce the reported findings are available in the Open Science Framework at https://osf.io/mjqwb/ (project identifier: mjqwb). The repository is organized by experiment (Exp1_Behavior, Exp2_TargetSelection_EEG, and Exp3_Visibility_Report_NoReport, the last with separate erp and mvpa folders); the source data for each figure and supplementary table are in the folder of the corresponding experiment.

## Abbreviations

2AFC: two-alternative forced-choice
AUC: area under the curve
EEG: electroencephalography
ERP: event-related potential
GLMM: generalized linear mixed model
ICA: independent component analysis
MVPA: multivariate pattern analysis
N2pc: N2-posterior-contralateral component
P3b: P3b component
ROI: region of interest
SOA: stimulus-onset asynchrony
VAN: visual awareness negativity.

## Supporting Information Captions

S1–S13 Figs and S1–S8 Tables are provided in the Supporting Information file.

## Supporting Information

**S1 Fig.**
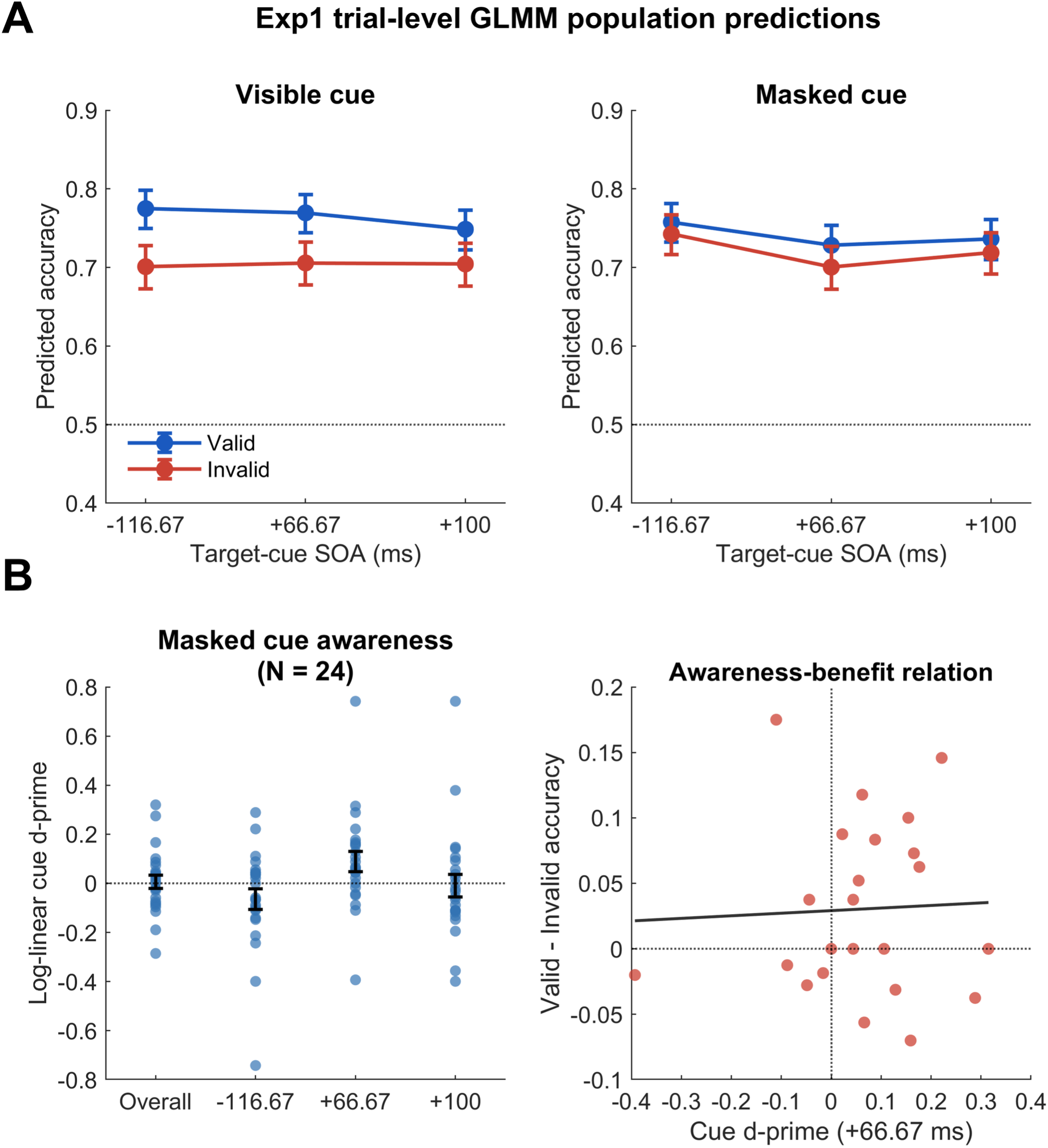
Experiment 1 behavioral, GLMM, and cue-awareness controls, related to Fig 2. A, Trial-level GLMM population predictions across SOAs (−116.67, +66.67, and +100 ms) for Valid and Invalid trials; points and intervals depict model predictions and 95% confidence intervals; the random-intercept model tier was selected by convergence and singularity diagnostics. B, Masked cue awareness, log-linear d-prime, and awareness-benefit relation across participants. Left/middle panels illustrate masked cue-location awareness performance and individual log-linear d-prime values; right panel shows the individual awareness-benefit correlation. The correlation is descriptive and is not interpreted as evidence for the complete absence of individual awareness.

**S2 Fig.**
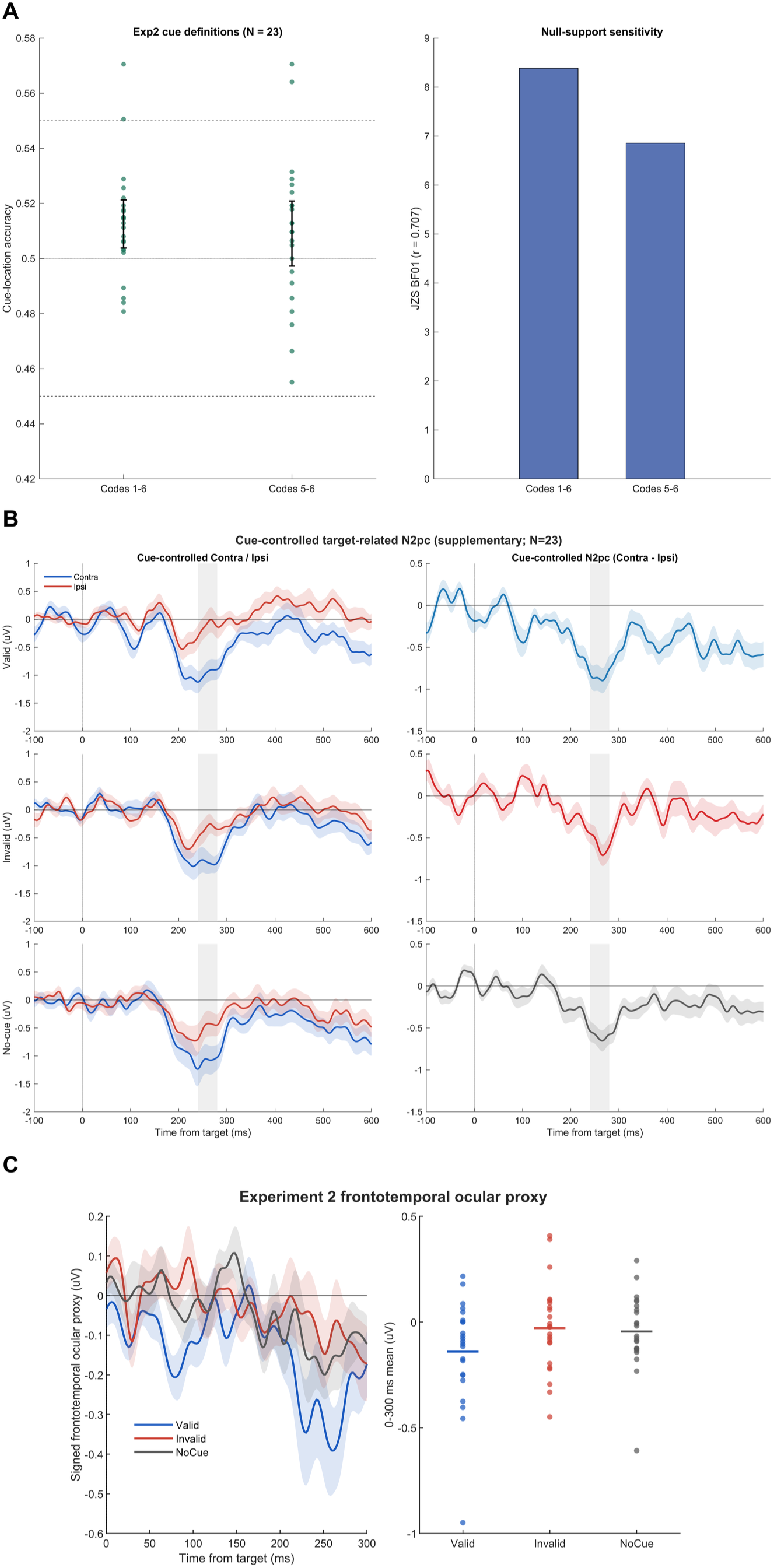
Experiment 2 cue-awareness definitions, cue-controlled target-related N2pc, and ocular controls, related to Fig 2 and S13 Fig. A, Experiment 2 cue-awareness definitions and bounds. Proportion of cue-present trials perceived across awareness code definitions; codes 1–6 denote the primary all-cue-present measure, while codes 5–6 denote the cue-only sensitivity definition. Dashed lines mark the ±5 percentage-point robustness bounds (±2.5 and ±7.5 percentage-point bounds are reported in S1 Table D). B, Experiment 2 cue-controlled target-related N2pc. Rows show Valid, Invalid, and No-cue conditions. Left, target-present minus cue-side-matched cue-only or mask-only Contra/Ipsi ROI waveforms after recoding relative to target side; right, the corresponding cue-controlled N2pc difference waves (Contra minus Ipsi). Lines show N = 23 group means and shading shows SEM; gray shading marks the pre-defined 240–280 ms window. This supplementary sensitivity analysis evaluates lateralized target-related activity after controlling cue-evoked contributions via cue-side-matched subtraction. C, Frontotemporal ocular proxy. The proxy is mean(Fp2,F8,FT8) minus mean(Fp1,F7,FT7), recoded as target-contralateral minus target-ipsilateral and averaged equally across target sides. No additional artifact-rejection threshold or post-processing was applied. The sub-plot beside A shows JZS Bayes factors (BF01) for each cue-awareness definition. The primary between-condition modulation was non-significant after cue control, whereas within-condition target-related lateralization remained reliable.

**S3 Fig.**
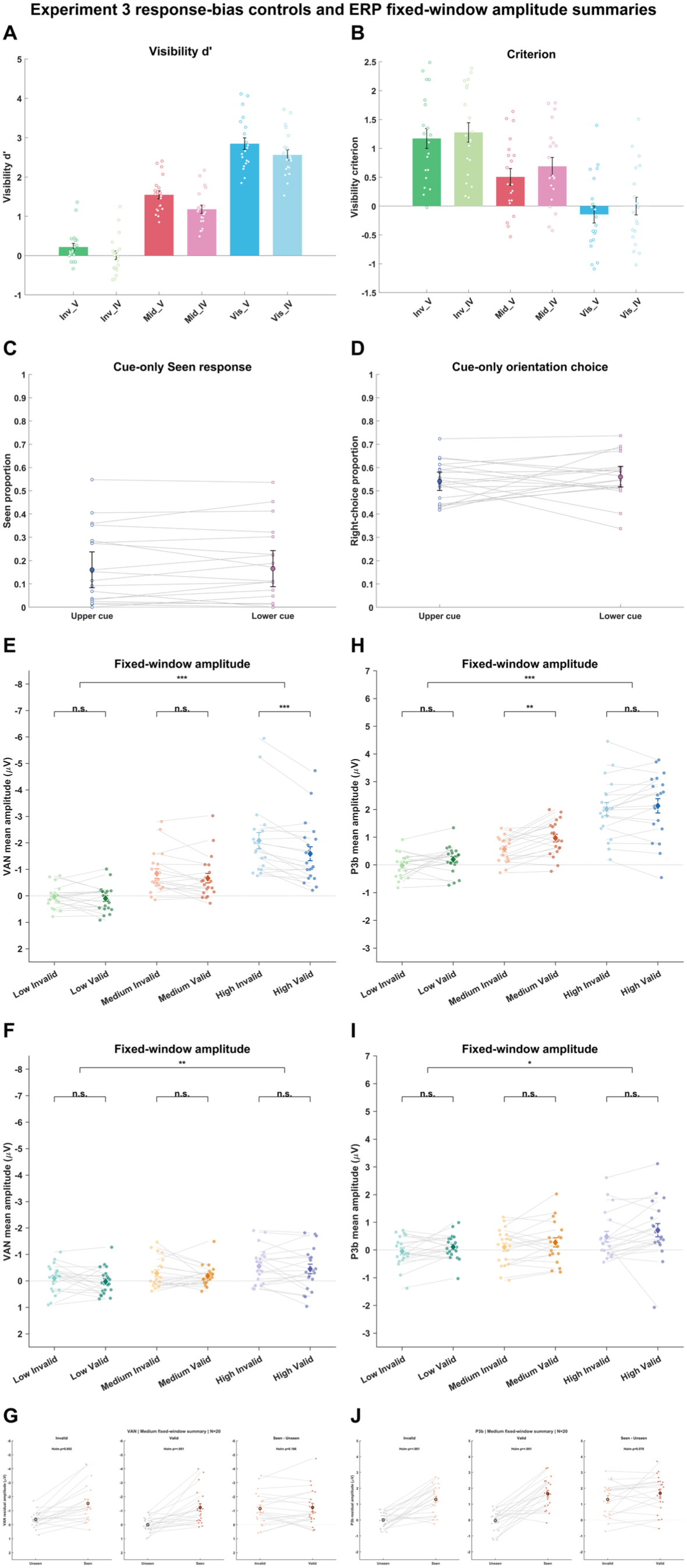
Experiment 3 response-bias controls and ERP fixed-window amplitude summaries, related to Figs 3 and 6. A–B, Experiment 3 visibility sensitivity (d′) and criterion estimates across target-present Valid and Invalid conditions. C–D, Cue-only catch-trial spatial response-bias controls. C, cue-only catch-trial Seen proportion following upper and lower cues. D, cue-only catch-trial right-tilted orientation-choice proportion following upper and lower cues. C– D display participant-level paired observations, group means, and 95% confidence intervals. Catch trials contained no target and therefore had no objective Valid/Invalid status. E–G, Visual Awareness Negativity (VAN) fixed-window (240–280 ms) amplitude summaries: E, Report-session amplitudes across Low, Medium, and High contrast for Invalid and Valid trials; F, corresponding No-report-session amplitudes; G, Medium-contrast Report-session VAN fixed-window Seen/Unseen summary within Invalid and Valid trials and the participant-level Seen-minus-Unseen comparison. H–J, Late positive complex (P3b) fixed-window (300–600 ms) amplitude summaries: H, Report-session amplitudes; I, No-report-session amplitudes; J, Medium-contrast Seen/Unseen summary within Invalid and Valid trials and the participant-level Seen-minus-Unseen comparison. Points show participants; large markers denote group means ± SEM; displayed tests are Holm-corrected.

**S4 Fig.**
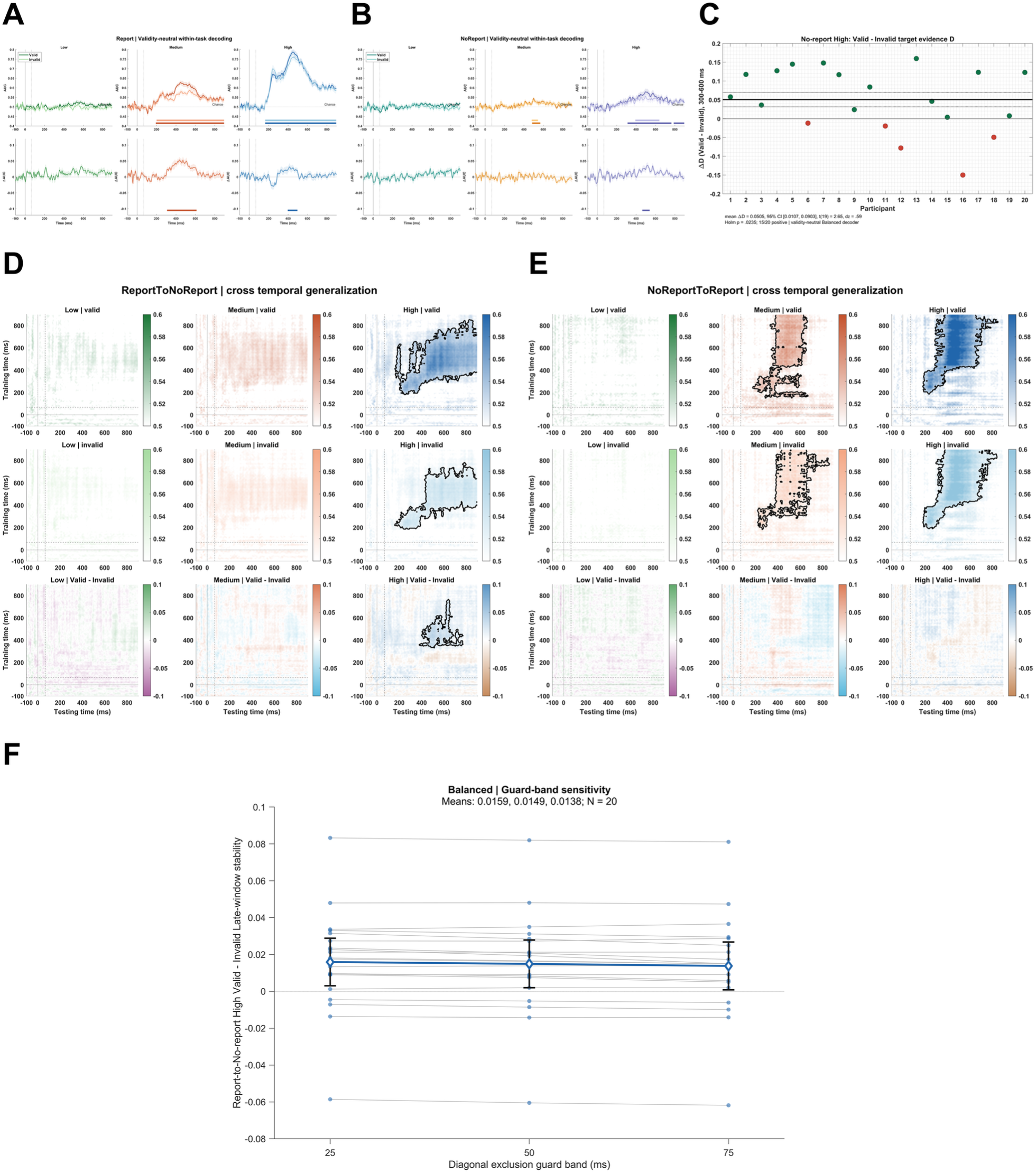
Primary Balanced decoder within-task performance and cross-task temporal generalization, related to Figs 4 and 5 and S8 Fig. A, Report-session target-versus-cue-only AUC time courses from the validity-neutral Balanced decoder for Valid and Invalid trials and their Valid-minus-Invalid differences across Low, Medium, and High contrast. B, Corresponding No-report-session target AUC time courses and paired differences. Horizontal bars denote corrected significance intervals. C, Late-window (300–600 ms) High-contrast No-report Valid-minus-Invalid target-evidence D summary for all 20 participants, with group mean, 95% confidence interval, and test statistics (15/20 participants positive). D, Balanced Report-to-No-report cross-task temporal generalization. Rows show Valid, Invalid, and Valid-minus-Invalid matrices across Low, Medium, and High contrast; black contours mark cluster-corrected significant clusters. E, Balanced No-report-to-Report reverse-direction temporal generalization, plotted with identical contrast-specific colormaps and complementary difference scales. F, Balanced late-window stability guard-band sensitivity. High-contrast Report-to-No-report Valid-minus-Invalid late-window stability across 25-, 50-, and 75-ms diagonal exclusions, displaying participant values, group means, and 95% confidence intervals.

**S5 Fig.**
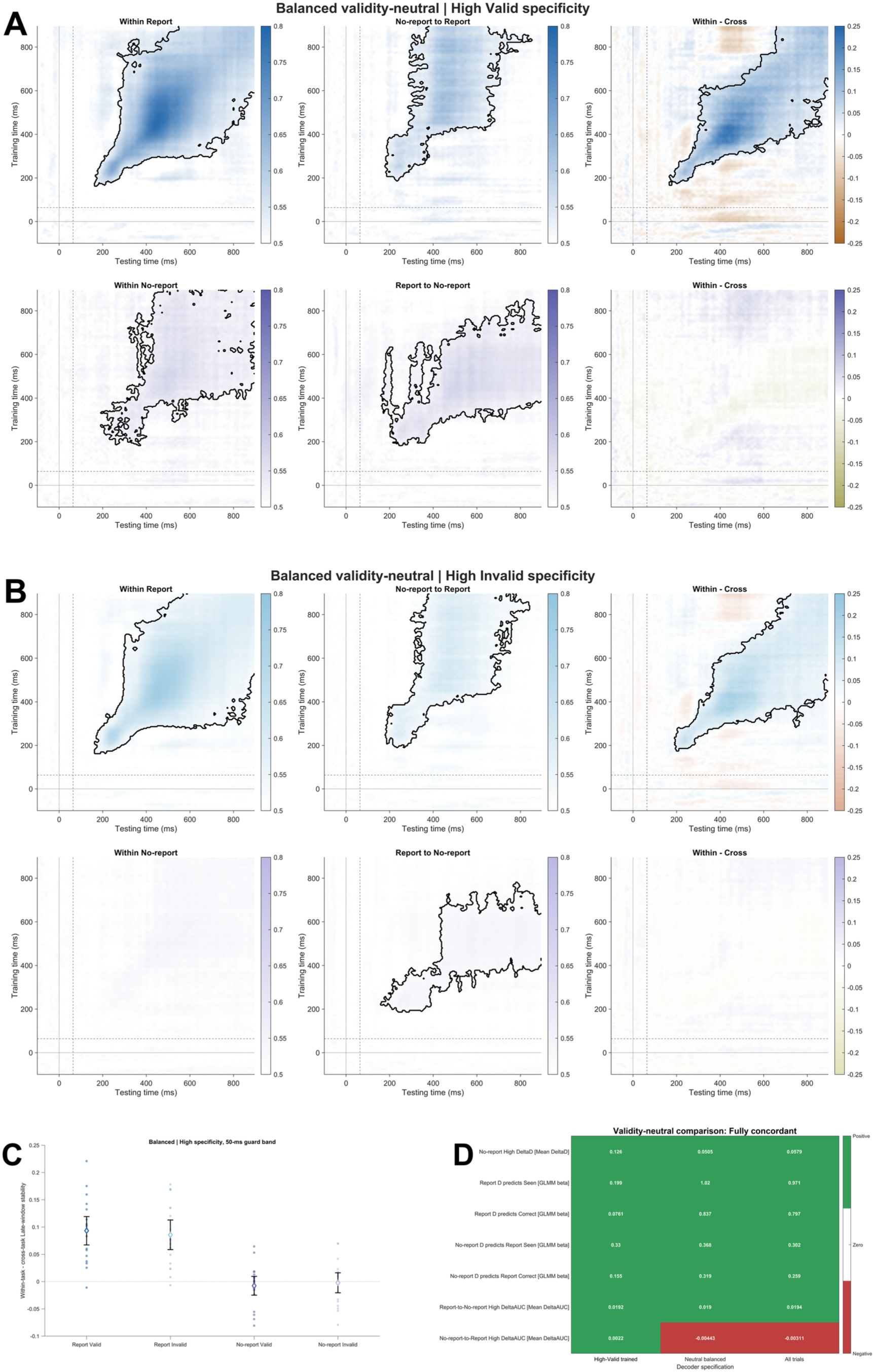
Primary Balanced task specificity and validity-neutral decoder comparison, related to Fig 5. A, Balanced High Valid task specificity. Within-task AUC, cross-task AUC, and within-minus-cross specificity matrices for each fixed test task (Report and No-report). Difference maps use a complementary negative-to-white-to-positive scale. B, Balanced High Invalid task specificity. Corresponding High Invalid specificity maps for the primary Balanced decoder. C, Balanced High specificity participant-level summary. Participant-level High-contrast late-window stability specificity scores with group means and 95% confidence intervals. (High-contrast specificity maps are also shown in Fig 5C–D.) D, Validity-neutral decoder comparison. Core outcome comparison across the High-Valid decoder, the primary Balanced decoder, and the all-trials sampling sensitivity, showing that the conclusions do not depend on the training approach.

**S6 Fig.**
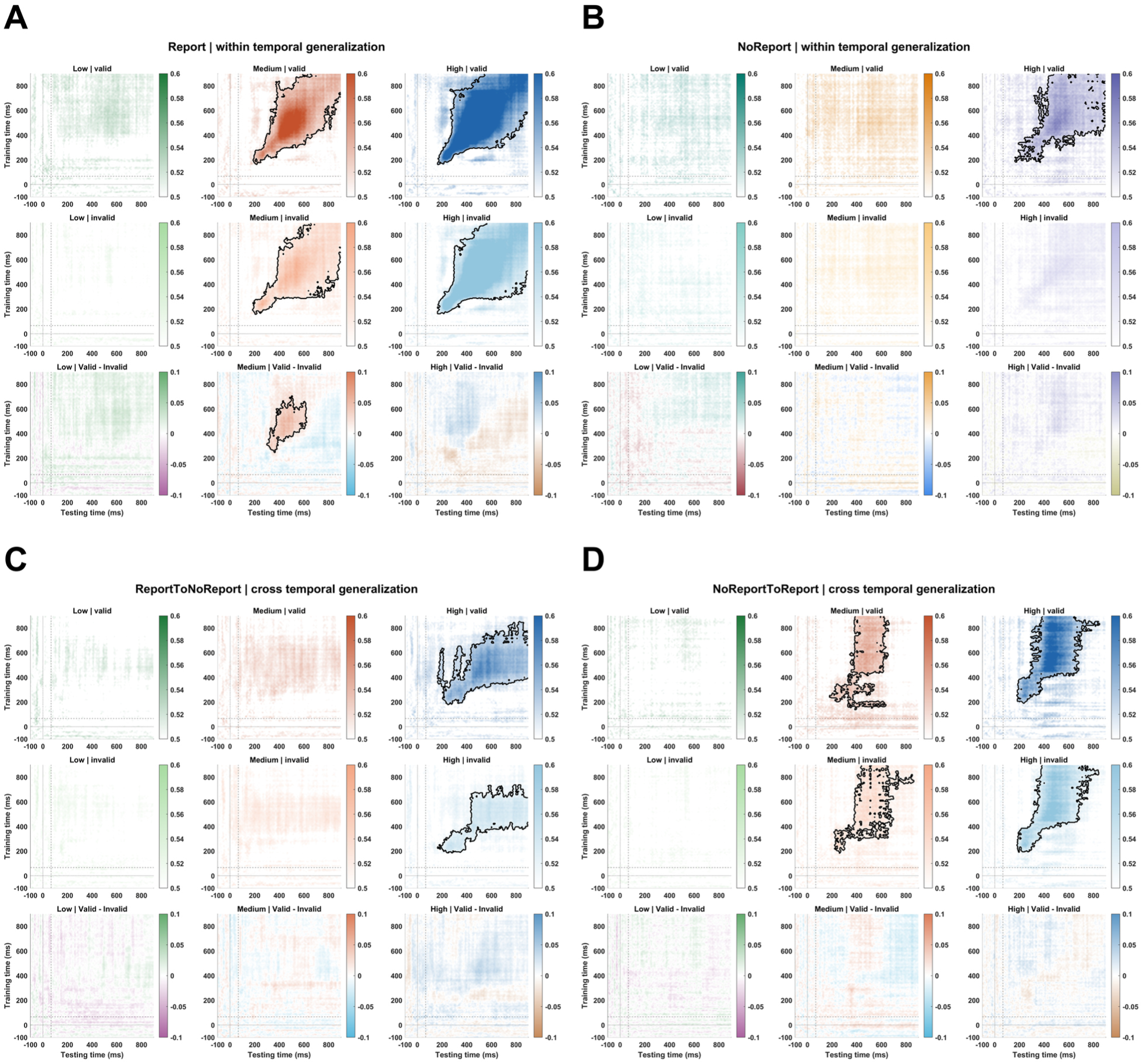
All-trials sampling sensitivity temporal generalization, related to Fig 5 and S8 Fig. All-trials sensitivity decoder trained across all available trials (sampling sensitivity check). A, Full within-Report temporal-generalization matrices across Low, Medium, and High contrast. B, Full within-No-report temporal-generalization matrices. C, Cross-task Report-to-No-report temporal-generalization matrices, plotted with identical contrast-specific colormaps as the primary decoder. D, Cross-task No-report-to-Report reverse-direction temporal-generalization matrices. Black contours denote corrected matrix clusters.

**S7 Fig.**
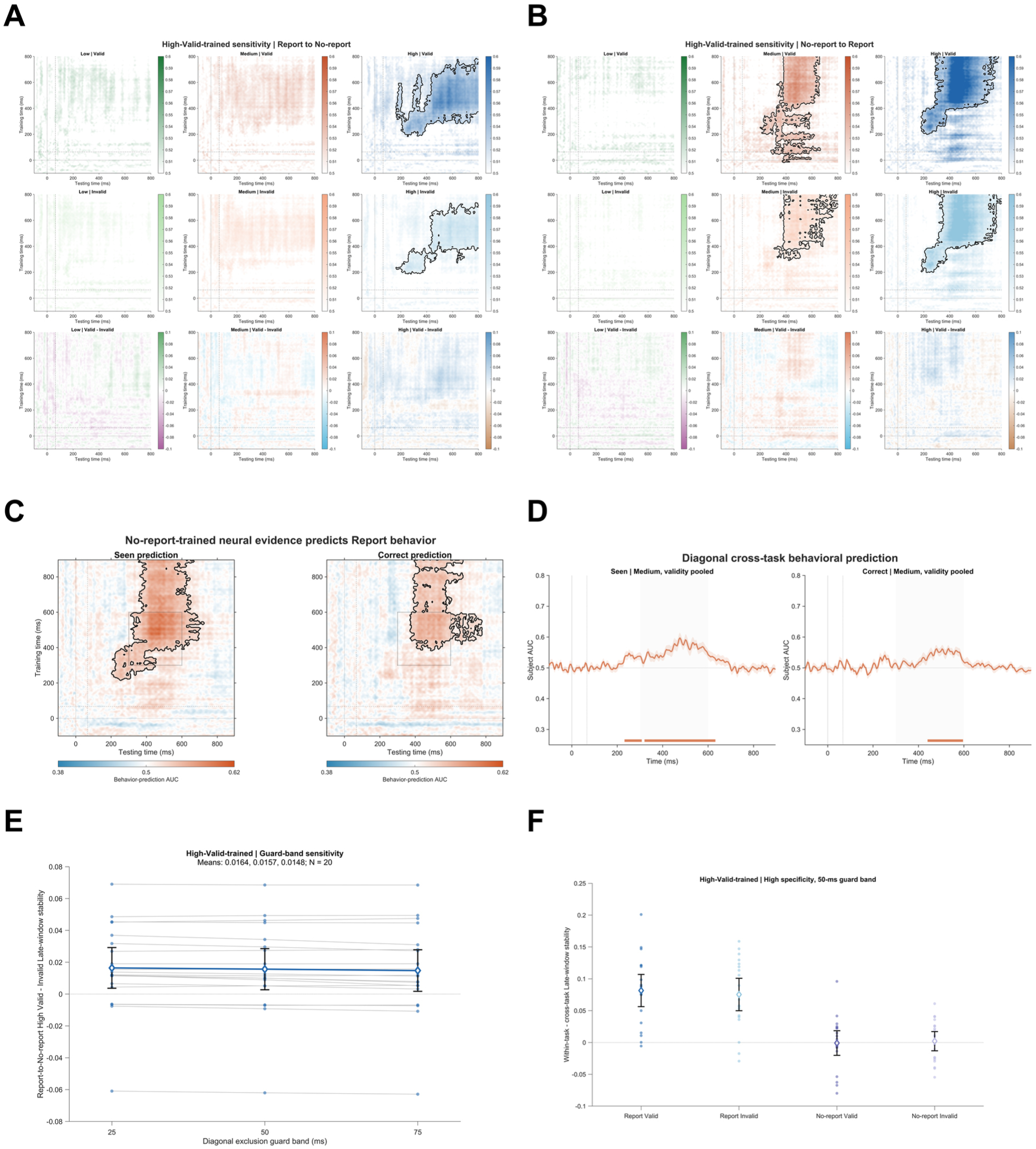
High-Valid-trained model neural and behavioral sensitivity analyses, related to Figs 4 and 5 and S8 Fig. Sensitivity analyses conducted with the alternative High-Valid-trained decoder model. A, High-Valid Report-to-No-report cross-task temporal generalization. B, High-Valid No-report-to-Report reverse-direction temporal generalization. C, High-Valid behavioral temporal generalization: No-report-trained neural evidence predicting independent Report Seen and Correct choices; black contours mark corrected clusters. D, High-Valid behavioral diagonal AUC time courses with formal significant intervals. E, High-Valid late-window stability guard-band sensitivity across 25-, 50-, and 75-ms diagonal exclusions. F, High-Valid participant-level specificity summary. (Secondary validity-stratified subpanels and numerical inventories are tabulated in S5 Table).

**S8 Fig.**
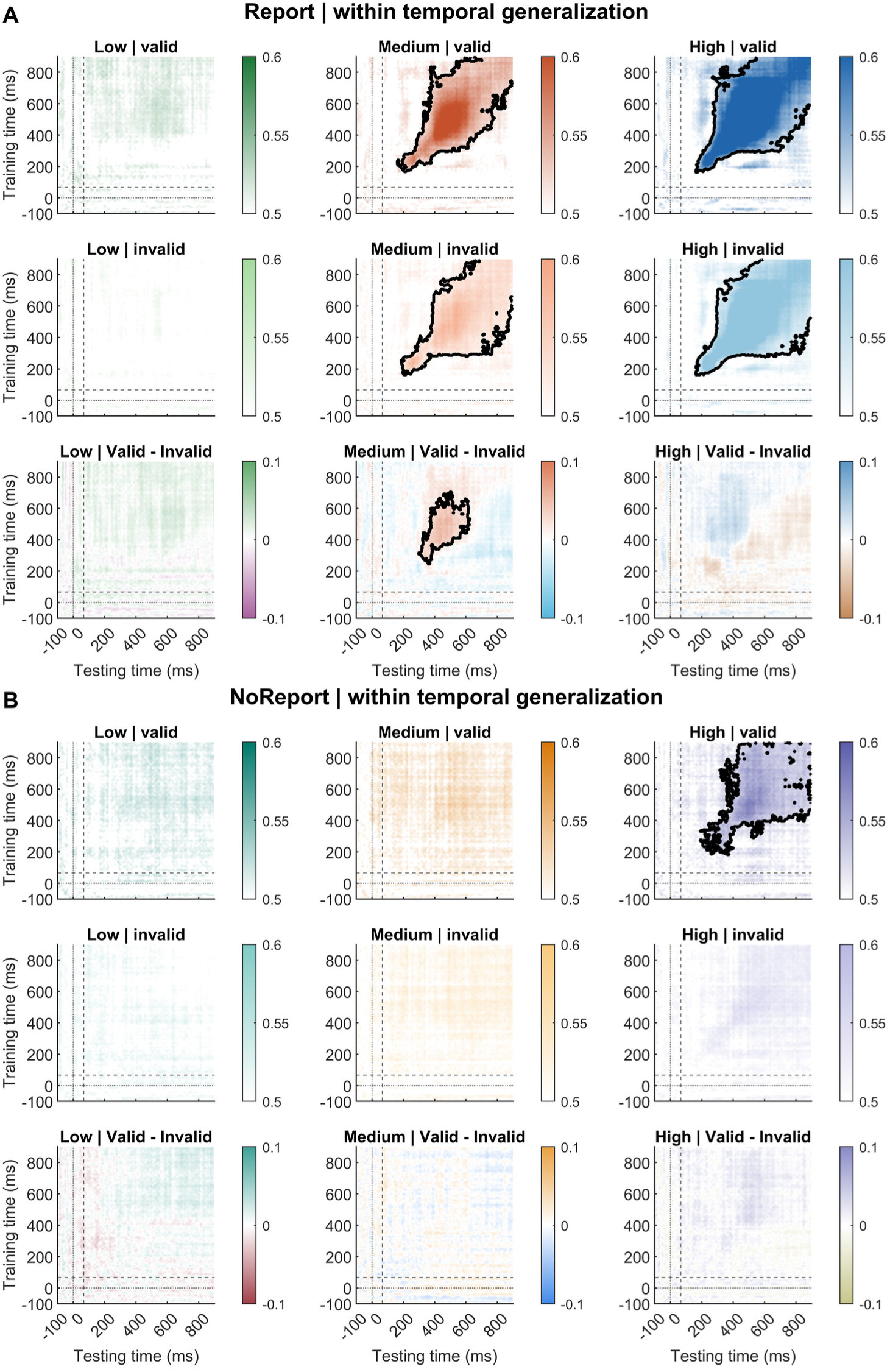
Within-task temporal generalization, related to Figs 4 and 5. Temporal generalization for Report (A) and No-report (B) from the validity-neutral Balanced decoder. Rows show Valid, Invalid, and Valid-minus-Invalid; columns show Low, Medium, and High contrast. Black contours denote corrected clusters. During Report, Medium-contrast Valid–Invalid temporal generalization formed a corrected 2D cluster spanning training 248–704 ms and testing 300–624 ms (cluster p = .0124; all-trials p = .0144). Under No-report, the High-contrast Valid–Invalid late-window stability index was positive (Balanced Δ = .00949, Holm-adjusted p = .444) without a corrected Valid–Invalid 2D cluster.

**S9 Fig.**
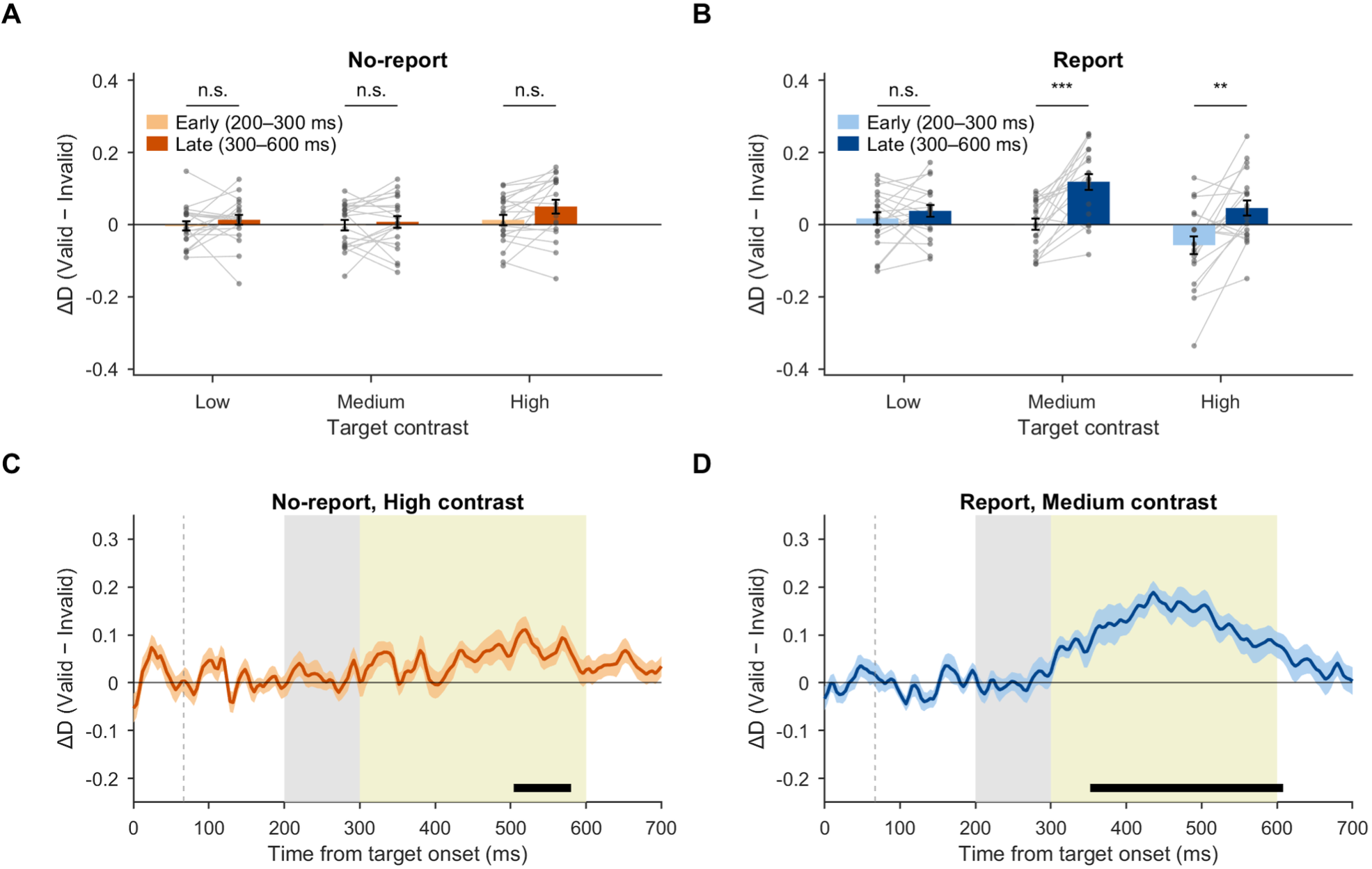
Late emergence of the cueing effect, related to Fig 4. A–B, ΔD (Valid − Invalid) in the early (200–300 ms) and late (300–600 ms) windows for each target contrast in No-report (A) and Report (B); bars show mean ± SEM and lines connect participants (N = 20); asterisks denote Holm-corrected paired tests of late versus early ΔD (family of six). C–D, time courses of ΔD for No-report High (C) and Report Medium (D) targets (mean ± SEM); shading marks the early and late windows, the dashed line marks cue onset (66.7 ms), and black bars mark corrected clusters (5,000 permutations).

**S10 Fig.**
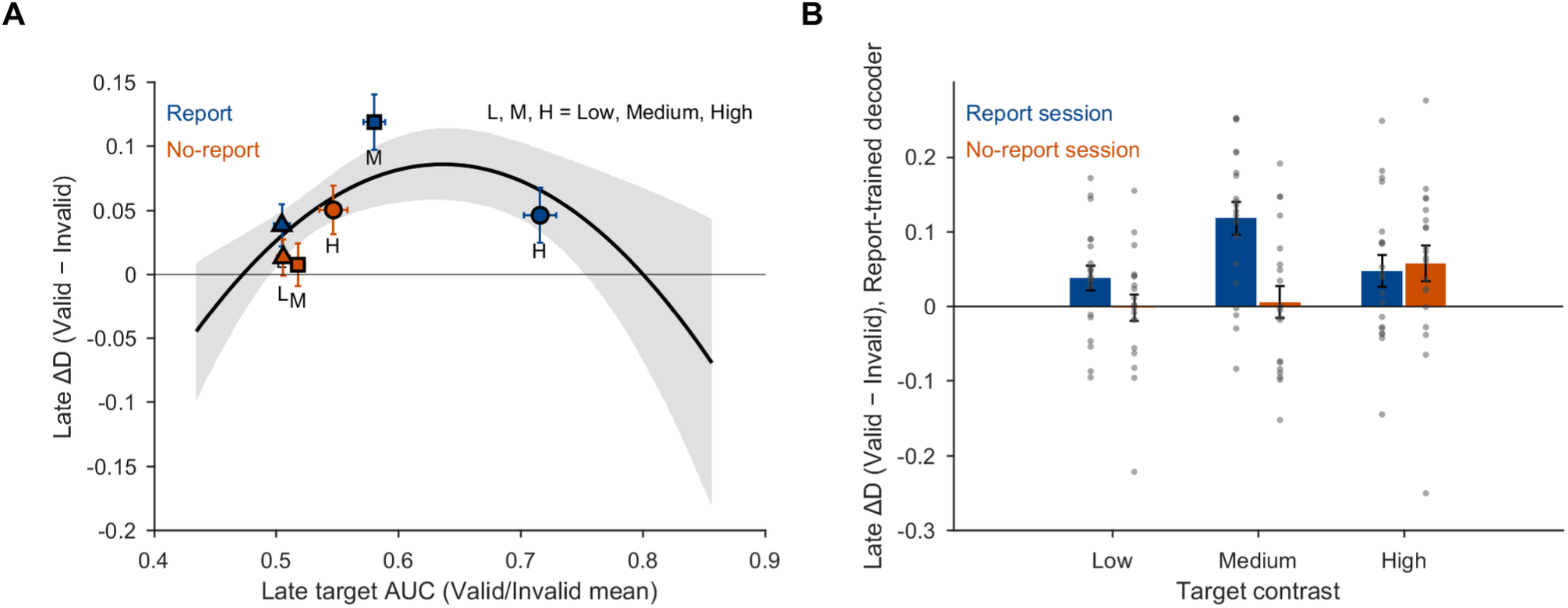
Target evidence and the cueing effect, related to Figs 4 and 5. A, late ΔD as a function of late target decodability (AUC) for the six task × contrast cells (mean ± SEM; blue, Report; orange, No-report), with the mixed-model quadratic fit and its 95% CI. B, late ΔD obtained by applying the Report-trained decoder to both sessions (mean ± SEM; dots show participants).

**S11 Fig.**
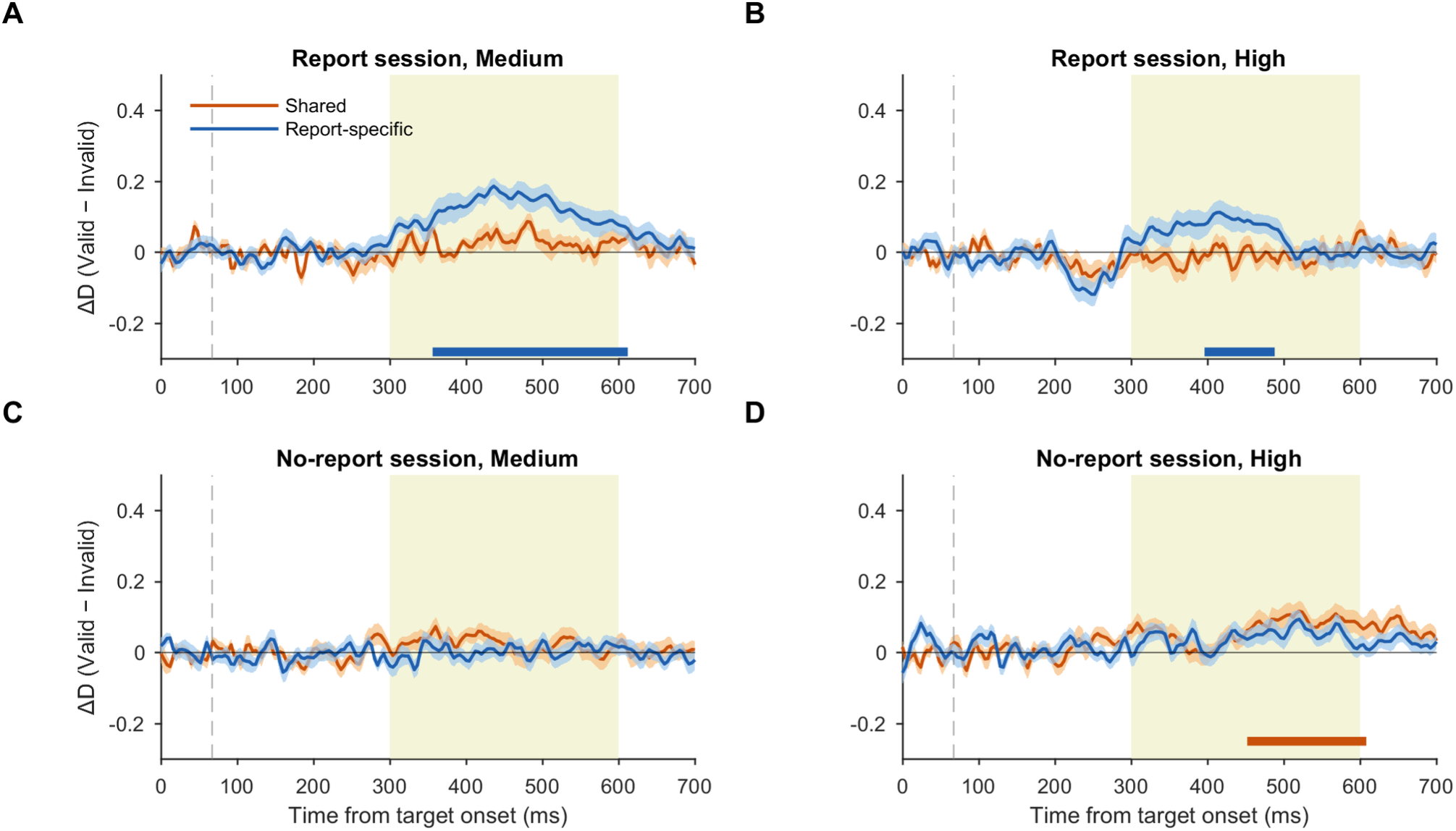
Time courses of the cueing effect on the shared and session-specific decoder components, related to Fig 5. ΔD (Valid − Invalid) on each component for Medium and High targets in the Report (A–B) and No-report (C–D) sessions (mean ± SEM; N = 20). In C–D, the shared component is the part of the No-report decoder along the Report decoder axis, and the specific component (blue) is orthogonal to it; horizontal bars mark corrected clusters (5,000 permutations; family of four per session).

**S12 Fig.**
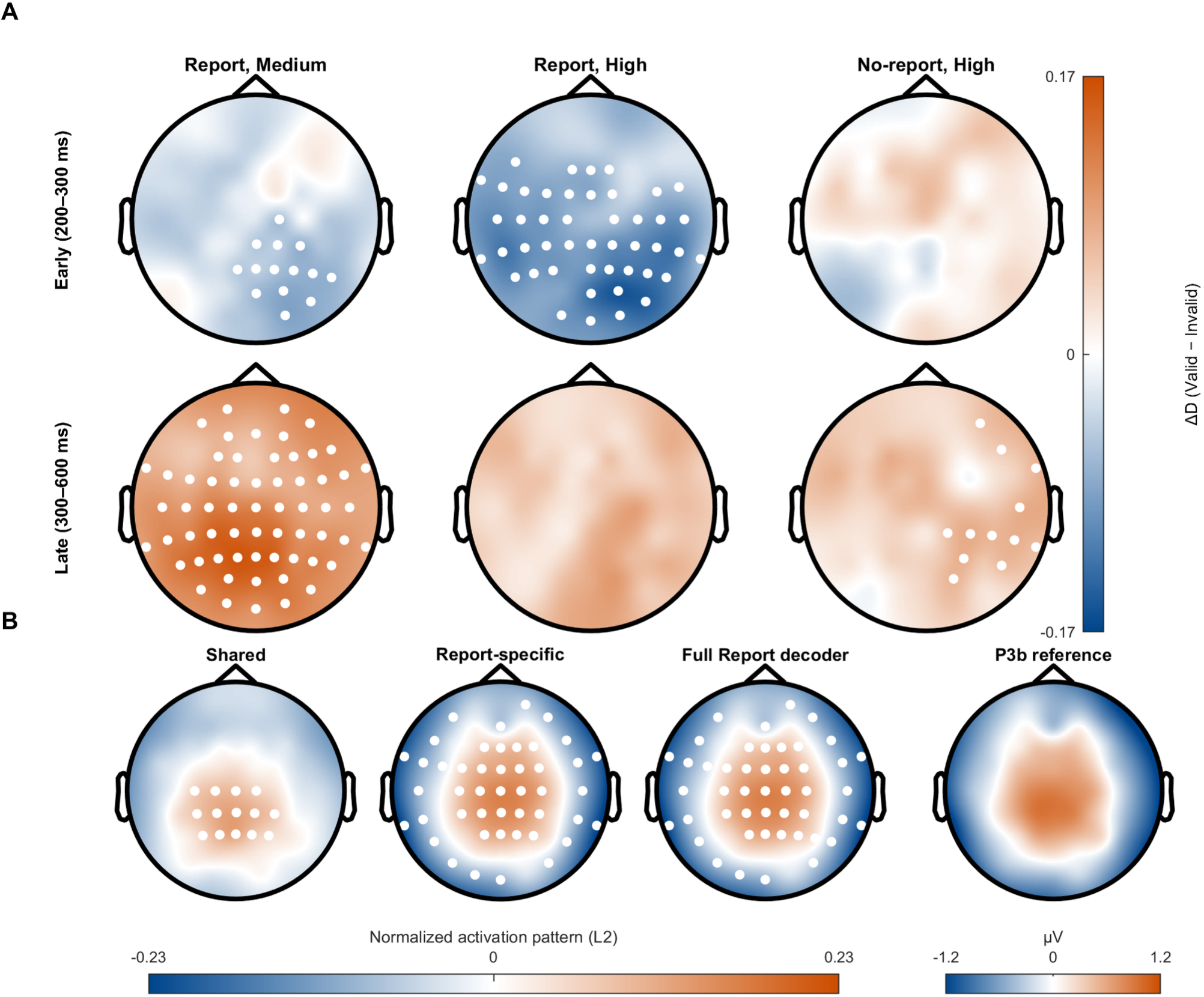
Scalp distribution of the cueing effect and activation patterns of the decoder components, related to Figs 4 and 5 and S9 Fig. A, searchlight ΔD (Valid − Invalid) maps in the early (200–300 ms) and late (300–600 ms) windows for Report Medium, Report High, and No-report High targets; white dots mark electrodes in significant clusters (cluster-based sign-flip permutations, 5,000; also in B). B, group-mean normalized activation patterns of the shared, Report-specific, and full Report decoders and the Report-session P3b reference topography (300–600 ms, Medium contrast). Topographies use the FieldTrip actiCAP 64-channel layout, as in Fig 6 (N = 20).

**S13 Fig.**
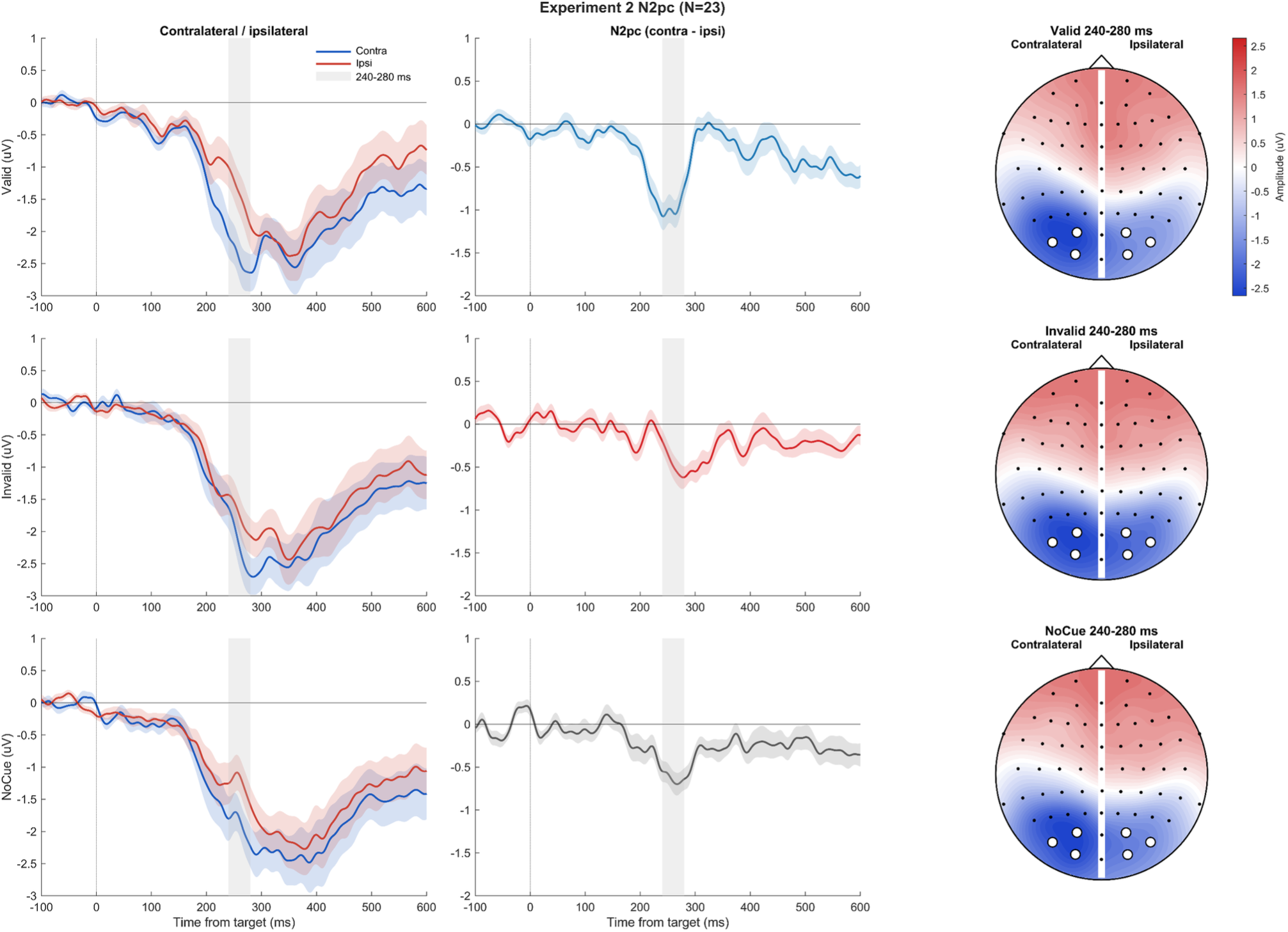
Experiment 2 N2pc, related to Fig 2. Rows show Valid, Invalid, and No-cue trials. Left, contralateral and ipsilateral ROI waveforms; middle, N2pc difference waves (contralateral minus ipsilateral), with the 240–280 ms window shaded; right, split-head scalp maps showing contralateral (left) and ipsilateral (right) activity; white circles mark O1/O2, PO3/PO4, and PO7/PO8. Positive voltage is plotted upward. N2pc differed across conditions (F(2,44) = 7.00, p = .0023), and Valid trials elicited a larger N2pc than Invalid (−0.519 µV, Holm-adjusted p = .0123) and No-cue trials (−0.344 µV, Holm-adjusted p = .0447). Cue-side-matched control analyses are shown in S2 Fig and S2 Table.

S1 Table. Experiment 1 behavioral, GLMM, and cue-awareness analyses, related to Fig 2.

S1 Table Notes. Cue-awareness inference. Cue-awareness was summarized at the participant level from trial-level cue-location accuracy. For each endpoint, a two-sided one-sample t test compared mean accuracy with 50%. A standard two-sided JZS point-null Bayes factor used H0: mean accuracy = 50% and H1: a non-zero standardized mean deviation with a Cauchy(0,r) prior, r = 1/√2 (0.7071); BF10 (H1/H0) and reciprocal BF01 (H0/H1) are reported. Independent two one-sided tests (TOSTs) evaluated ±2.5, ±5, and ±7.5 percentage-point bounds at alpha = 0.05. These are robustness bounds; the ±5 percentage-point (45–55%) bound is a percentage-point equivalence bound defined for the present robustness analysis rather than a preregistered criterion.

Experiment 1 inclusion sensitivity. All 29 recruited participants were included in a sensitivity refit while the 65–85% rule defined the primary N = 24 cohort. The sensitivity refit used the same Correct ∼ Validity * SOA * CueVisibility fixed-effects formula, effect coding, Laplace fitting, random-effects tier selection, Type-III Wald tests, and four-term Holm correction. Participant-level masked Valid-Invalid contrasts at +66.67 ms and overall were analyzed as paired sensitivity checks. The GLMM pattern was retained in the all-recruited refit: Validity remained significant (F(1,25741) = 47.22, Holm p = 2.60 × 10⁻¹¹) and Validity × Cue visibility remained significant (F(1,25741) = 16.94, Holm p = 1.16 × 10⁻⁴), whereas Validity × SOA (F(1,25741) = 2.30, Holm p = 0.2588) and the three-way term (F(1,25741) = 0.625, Holm p = 0.4292) remained non-significant, matching the N = 24 primary results (S1 Table C). In contrast, the key cell-level retrospective benefit at +66.67 ms did not replicate across the full sample: the participant-level masked Valid-minus-Invalid paired contrast was t(22) = 2.203, raw p = 0.0384 (mean = +3.04 percentage points) in the retained N = 24 cohort but t(27) = -0.114, raw p = 0.910 (mean = -0.26 percentage points) in the all-recruited N = 29 sensitivity. The masked +66.67-ms benefit therefore depends on the 65–85% overall-accuracy inclusion criterion; the N = 24 cohort is reported as primary, and the all-recruited analysis is reported here as a sensitivity analysis. The paired +66.67-ms sensitivity contrasts used pairwise-complete participants (n = 23 in the retained cohort and n = 28 in the all-recruited cohort); the excluded records lacked an estimable Valid/Invalid pair.

S2 Table. Experiment 2 behavioral, cue-awareness, EEG, and QC controls, related to Fig 2 and S13 Fig.

S2 Table Notes. Experiment 2 behavioral and EEG control analyses. Experiment 2 behavior was summarized at the participant level from the retained blocks. All cue-present accuracy definitions were computed at the participant level. EEG epochs were aligned to target onset (64 channels, 500 Hz, after ICA); behavioral response and correctness labels were not attached to EEG epochs. N2pc used O1/O2, PO3/PO4, and PO7/PO8 at 240–280 ms with equal left/right target weighting.

Experiment 2 behavior. Across 23 participants, pooled orientation accuracy was 75.00% for Valid, 72.91% for Invalid, and 73.85% for No-cue. The condition effect was not significant, F(2,44) = 2.69, p = .0794. Valid-Invalid was +2.09 percentage points (t(22) = 2.88, raw p = .00863, Holm p = .0259). The primary all-cue-present cue-location definition (codes 1–6, row 10) was 51.25% (BF10 = 6.75, BF01 = 0.15); the cue-only codes 5–6 sensitivity was 50.91% (BF10 = 0.66, BF01 = 1.53). The ±2.5, ±5, and ±7.5 percentage-point TOST bounds were met for both definitions (S1 Table D). The all-cue estimate is described as small residual discrimination, not exact chance.

Experiment 2 N2pc. Mean contralateral-minus-ipsilateral amplitudes were -0.977, -0.458, and -0.633 µV for Valid, Invalid, and No-cue. The primary difference-wave condition effect was F(2,44) = 7.00, p = .00230; Valid was more negative than Invalid and No-cue. In the cue-controlled sensitivity analysis, the condition effect was attenuated, F(2,44) = 2.60, GG p = .0928, and the two planned Valid contrasts had Holm p = .1991. The frontotemporal ocular proxy itself showed no condition effect from 0–300 ms, F(2,44) = 2.54, p = .0906. Adding the proxy as a covariate yielded adjusted Valid-Invalid and Valid-No-cue contrasts of -0.375 and -0.264 µV, respectively.

S3 Table. Experiment 3 behavioral and response-bias analyses, related to Fig 3.

S3 Table Notes. Cue-only catch-trial spatial response-bias control. Cue-only catch trials contained no Gabor target and therefore had no target location or Valid/Invalid status. We analyzed Report-session upper-cue and lower-cue catch trials with valid four-option responses. For each participant, upper- and lower-cue Seen proportions were compared with a two-sided paired t test. Orientation response was coded as left-tilted for response bins 1/2 and right-tilted for bins 3/4 and tested at trial level with the binomial-logit mixed model RightChoice ∼ CueUpper + (1 + CueUpper | participant), fitted by Laplace approximation. The two planned cue-location tests were Holm-corrected.

Seen-response proportions were 16.05% after upper cues and 16.56% after lower cues (upper-minus-lower mean difference = −0.51 percentage points, 95% CI [−2.95, 1.92], t(19) = −0.44, raw p = .664, Holm-adjusted p = .888). Right-tilted orientation choices occurred on 54.13% of upper-cue and 56.05% of lower-cue catch trials. The trial-level mixed model gave β = −0.075, 95% CI [−0.266, 0.117], z = −0.77, OR = 0.928, 95% CI [0.766, 1.124], raw p = .444, and Holm-adjusted p = .888. Neither outcome showed evidence of a cue-location-induced response bias. This analysis tests spatial bias induced by cue location itself; it is not a Valid/Invalid comparison and cannot by itself exclude all validity-related criterion shifts on target-present trials.

S4 Table. Primary Balanced decoder inference and temporal generalization, related to Figs 4 and 5 and S8 Fig.

S4 Table is provided as an Excel file. The workbook comprises eight worksheets corresponding to the primary validity-neutral Balanced decoding analyses: S4 Table A. Balanced corrected cluster summary; S4 Table B. Primary Balanced 50-ms late-window stability tests; S4 Table C. Report-to-No-report High guard-band sensitivity; S4 Table D. Primary one-dimensional neural-evidence and behavior-prediction clusters; S4 Table E. Balanced High-contrast task specificity: corrected matrix clusters; S4 Table F. Balanced High-contrast late-window stability specificity tests; S4 Table G. Balanced specificity repeated-measures model; S4 Table H. Balanced and all-trials sensitivity summary.

Analysis overview. The validity-neutral Balanced decoder (primary) is the inferential basis for the main text and primary supplementary results. The all-trials decoder (sampling sensitivity) tests whether equal-midpoint use of all eligible trials changes conclusions. The High-Valid-trained decoder serves only as a sensitivity analysis.

The primary decoder was trained independently of cue validity. Subject-level out-of-fold evidence was used for behavior prediction, and source-task training was kept separate from independent target-task testing. All inferential summaries below use participants as the statistical unit (N = 20 unless a table states otherwise). Cluster procedures used 5,000 participant-level sign flips with the predefined correction family. For one-dimensional held-out D, Report and No-report used the same procedure: condition-wise D-versus-zero tests formed a six-curve family (Low/Medium/High by Valid/Invalid; right-tailed), whereas Valid-minus-Invalid ΔD tests formed a separate three-contrast family (two-sided); both used the same max-cluster correction procedure.

Temporal-generalization matrices cover the full analysed epoch. Black contours denote corrected clusters. Valid-minus-Invalid maps use high-contrast complementary negative and positive colors around a white zero. Late-window stability is the mean Valid-minus-Invalid AUC in the 300–600-ms train-by-test region after excluding a diagonal guard band; 50 ms is primary and 25/75 ms are descriptive sensitivity checks. Task specificity was evaluated while holding the test task fixed: the corresponding cross-task temporal-generalization matrix was subtracted from the within-task matrix separately for Valid and Invalid trials. Positive within-minus-cross values therefore index target-related information expressed more strongly when training and testing occurred within the same task than could be transferred from the other task. The four High-contrast specificity maps (Report-Valid, Report-Invalid, No-report-Valid, and No-report-Invalid) formed a single four-panel two-sided 2D cluster-permutation family with 5,000 permutations, a max-sum cluster statistic, and minimal adjacency; positive and negative clusters were each tested at α/2 within the shared max-cluster family. Participant-level specificity late-window stability was the mean within-minus-cross AUC in the 300–600-ms train-by-test region after excluding |train−test| ≤ 50 ms; the four High-contrast task-by-validity tests were two-sided and Holm-corrected together. Selected High-contrast specificity difference maps are foregrounded in Fig 5C–D of the main text, while the full component matrices and participant-level summaries are shown in S5 Fig.

For the primary Balanced decoder, Report-trained High-contrast Valid-minus-Invalid evidence generalized to No-report in a corrected cross-temporal cluster spanning training 332–764 ms and testing 380–768 ms (p = .0234). The corresponding Report-to-No-report High late-window stability means were .0159, .0149, and .0138 for 25-, 50-, and 75-ms guard bands, respectively, with 16/20 participants positive at the primary 50-ms setting. These results are based on the primary validity-neutral Balanced decoder; High-Valid-trained results are reported separately as sensitivity analyses.

High-contrast task specificity for the Balanced decoder was positive when Report was the fixed test task: mean within-minus-cross differences were .0932 and .0858 for Valid and Invalid trials. Corrected clusters covered training/testing 168–896 ms for Valid and training 176–896 ms/testing 192–896 ms for Invalid (both p = .0002). No-report-test specificity did not show the corresponding positive effect.

The primary one-dimensional neural-evidence and behavior-prediction results are concordant with the plotted significance bars. In Report, Medium Valid-minus-Invalid ΔD was significant from 352–608 ms (p = .0002), whereas High ΔD had no corrected cluster (Fig 4A). In No-report, High Valid-minus-Invalid ΔD remained significant from 504–580 ms (p = .0054; Fig 4B). The complete condition-wise D, ΔD, and behavior-prediction cluster inventory is given in S4 Table D. No preselected 300–600-ms shading is used as a substitute for the 0–900-ms cluster-permutation results.

S5 Table. Decoder robustness, sensitivity, and leakage controls, related to Figs 4 and 5 and S8 Fig.

S5 Table Notes. Balanced and all-trials validity-neutral models address the same design with different eligible-trial sampling. High-Valid neural and behavioral analyses are sensitivity comparisons and are not used to redefine the primary decoder or its inferential claims. Leakage control. Training and behavior projection were separated by participant and fold, and cross-task decoders were trained on the source task and evaluated on the independent target task. Checks confirmed that every training fold contained trials and that every trial received an out-of-fold value.

High-Valid behavioral temporal generalization is reported as a sensitivity analysis; its clusters were corrected as in the primary analysis.

S6 Table. Experiment 3 stimulus calibration, related to Materials and Methods.

S6 Table Notes. Experiment 3 stimulus calibration. Final participant-level contrast values were taken from the configuration values used for stimulus presentation. Report and No-report values are summarized separately because the session-specific saved parameters were not identical for all participants. The display calibration used a luminance-meter-derived mapping from integer RGB gray level to luminance in nit (cd/m^2). For each saved contrast c, the nominal dark and bright RGB endpoints were obtained by converting 0.2902 − c and 0.2902 + c to 8-bit gray levels; their calibrated luminances were used as Lmin and Lmax in C_M = (Lmax-Lmin)/(Lmax+Lmin). The calibration was applied post hoc to the saved RGB-domain stimulus parameters and was not used online as a Psychtoolbox gamma LUT.

As a descriptive check of the spacing of the three calibrated stimulus levels, the group-mean luminance-calibrated Michelson contrasts increased approximately linearly from Low to Medium to High. The regression used level index Low = 1, Medium = 2, and High = 3 as the predictor and the three session-level group means as the outcome, with Report and No-report fitted separately (N = 3 group-level contrast levels per fit). The R^2 values characterize spacing only and are not an inferential test.

S7 Table. Experiment 3 No-report engagement, logging, and provenance, related to Fig 3 and Materials and Methods.

S7 Table Notes. Across the cohort, the nominal experimental schedule was four 196-trial blocks (784 trials; 15,680 slots across 20 participants). In the EEG cohort (N = 20), 18 participants completed four 196-trial blocks, one participant completed four 182-trial blocks (728 trials), and one participant completed five blocks (including a 112-trial fifth block; 896 trials), yielding 15,736 trials in total (11,240 engagement events, 2,248 thought probes, and 2,248 continuation prompts).

Category proportions were summarized across participants with at least one valid thought-probe response (N = 18); missing/invalid response summaries include all participants (N = 20). There were 2,248 configured probes, of which 2,004 were valid and 244 were missing or invalid. Eighteen of 20 participants had at least one valid probe response. These probes are intermittent and indirect samples of ongoing conscious content and were not used to assign trial-wise visibility labels to Gabor targets.

Together, the sparse speed task, thought probes, and simple continuation prompt indicate that participants remained behaviorally responsive to orthogonal events across all 15,736 trials (18 participants with 784 trials, one with 728 trials, and one with 896 trials). The 104 unrecorded prompt slots for one participant are treated as missing records rather than behavioral noncompliance. These measures do not provide trial-wise Seen/Unseen or Gabor-awareness labels in the No-report session.

S8 Table. Supplementary decoder analyses, related to Figs 4 and 5 and S9–S11 Figs. Late-window cueing effects on the shared and Report-specific components (primary, pattern-based, and regression-based definitions), behavioral GLMMs for both components, target-evidence and common-axis models, searchlight cluster tests and anterior–posterior indices, activation-pattern statistics, direct comparisons of the two components, session-order analyses, the No-report cueing effect after excluding trials that followed a thought probe, and all Experiment 3 results recomputed after excluding the participant who was an author (N = 19).

## References

1. Sergent C, Wyart V, Babo-Rebelo M, Cohen L, Naccache L, Tallon-Baudry C. Cueing attention after the stimulus is gone can retrospectively trigger conscious perception. Curr Biol. 2013;23:150–155. doi: 10.1016/j.cub.2012.11.047.

2. Thibault L, van den Berg R, Cavanagh P, Sergent C. Retrospective Attention Gates Discrete Conscious Access to Past Sensory Stimuli. PLoS One. 2016;11:e0148504. doi: 10.1371/journal.pone.0148504.

3. Tsuchiya N, Wilke M, Frässle S, Lamme VAF. No-Report Paradigms: Extracting the True Neural Correlates of Consciousness. Trends Cogn Sci. 2015;19:757–770. doi: 10.1016/j.tics.2015.10.002.

4. Mulckhuyse M, Theeuwes J. Unconscious attentional orienting to exogenous cues: A review of the literature. Acta Psychol (Amst). 2010;134:299–309. doi: 10.1016/j.actpsy.2010.03.002.

5. Pitts MA, Lutsyshyna LA, Hillyard SA. The relationship between attention and consciousness: an expanded taxonomy and implications for ‘no-report’ paradigms. Philos Trans R Soc Lond B Biol Sci. 2018;373:20170348. doi: 10.1098/rstb.2017.0348.

6. Lamme VA. Why visual attention and awareness are different. Trends Cogn Sci. 2003;7:12–18.

7. Chica AB, Lasaponara S, Chanes L, Valero-Cabre A, Doricchi F, Lupianez J, et al. Spatial attention and conscious perception: the role of endogenous and exogenous orienting. Atten Percept Psychophys. 2011;73:1065–1081.

8. Dehaene S, Lau H, Kouider S. What is consciousness, and could machines have it? Science. 2017;358:486–492.

9. Dehaene S. Consciousness and the brain: Deciphering how the brain codes our thoughts. Viking; 2014.

10. Block N. Two neural correlates of consciousness. Trends Cogn Sci. 2005;9:46–52.

11. Aru J, Bachmann T, Singer W, Melloni L. Distilling the neural correlates of consciousness. Neurosci Biobehav Rev. 2012;36:737–746.

12. Cohen MA, Ortego K, Kyroudis A, Pitts M. Distinguishing the Neural Correlates of Perceptual Awareness and Postperceptual Processing. J Neurosci. 2020;40:4925–4935.

13. Giattino CM, Alam ZM, Woldorff MG. Neural processes underlying the orienting of attention without awareness. Cortex. 2018;102:14–25.

14. Prasad S, Mishra RK. The Nature of Unconscious Attention to Subliminal Cues. Vision (Basel). 2019;3:38.

15. Hatamimajoumerd E, Ratan Murty NA, Pitts M, Cohen MA. Decoding perceptual awareness across the brain with a no-report fMRI masking paradigm. Curr Biol. 2022;32:4139–4149.e4. doi: 10.1016/j.cub.2022.07.068.

16. Sergent C, Corazzol M, Labouret G, Stockart F, Wexler M, King J-R, et al. Bifurcation in brain dynamics reveals a signature of conscious processing independent of report. Nat Commun. 2021;12:1149. doi: 10.1038/s41467-021-21393-z.

17. Cohen MA, Dembski C, Ortego K, Steinhibler C, Pitts M. Neural signatures of visual awareness independent of postperceptual processing. Cereb Cortex. 2024;34:bhae415. doi: 10.1093/cercor/bhae415.

18. Kronemer SI, Aksen M, Ding JZ, Ryu JH, Xin Q, Ding Z, et al. Human visual consciousness involves large scale cortical and subcortical networks independent of task report and eye movement activity. Nat Commun. 2022;13:7342.

19. Dellert T, Muller-Bardorff M, Schlossmacher I, Pitts M, Hofmann D, Bruchmann M, et al. Dissociating the Neural Correlates of Consciousness and Task Relevance in Face Perception Using Simultaneous EEG-fMRI. J Neurosci. 2021;41:7864–7875.

20. Stockart F, Msheik R, Robin A, Jurkovičová L, Goueytes D, Rouy M, et al. Cortical evidence accumulation for visual perception occurs irrespective of reports. Nat Commun. 2025;16:8458. doi: 10.1038/s41467-025-63255-y.

21. Cogitate Consortium, Ferrante O, Gorska-Klimowska U, Henin S, Hirschhorn R, Khalaf A, et al. Adversarial testing of global neuronal workspace and integrated information theories of consciousness. Nature. 2025;642:133–142. doi: 10.1038/s41586-025-08888-1.

22. Rimsky-Robert D, Stormer V, Sackur J, Sergent C. Retrospective auditory cues can improve detection of near-threshold visual targets. Sci Rep. 2019;9:18966.

23. Xia Y, Morimoto Y, Noguchi Y. Retrospective triggering of conscious perception by an interstimulus interaction. J Vis. 2016;16:3.

24. Eimer M. The N2pc component as an indicator of attentional selectivity. Electroencephalogr Clin Neurophysiol. 1996;99:225–234. doi: 10.1016/0013-4694(96)95711-9.

25. Vogel EK, Woodman GF, Luck SJ. Pushing around the locus of selection: evidence for the flexible-selection hypothesis. J Cogn Neurosci. 2005;17:1907–1922.

26. Eklund R, Wiens S. Visual awareness negativity is an early neural correlate of awareness: A preregistered study with two Gabor sizes. Cogn Affect Behav Neurosci. 2018;18:176–188. doi: 10.3758/s13415-018-0562-z.

27. Del Cul A, Baillet S, Dehaene S. Brain Dynamics Underlying the Nonlinear Threshold for Access to Consciousness. PLoS Biol. 2007;5:e260.

28. King JR, Dehaene S. Characterizing the dynamics of mental representations: the temporal generalization method. Trends Cogn Sci. 2014;18:203–210. doi: 10.1016/j.tics.2014.01.002.

29. Pitts MA, Padwal J, Fennelly D, Martinez A, Hillyard SA. Gamma band activity and the P3 reflect post-perceptual processes, not visual awareness. Neuroimage. 2014;101:337–350.

30. Nicolacoudis A. Decoding EEG Bifurcation Dynamics Around Visual Detection Threshold in No-report. [thesis]. Reed College; 2025.

31. Baars BJ. *Global workspace theory of consciousness: toward a cognitive neuroscience of human experience*. In: Laureys S, editor. Progress in Brain Research. Elsevier; 2005. p. 45–53.

32. Dehaene S, Changeux JP. Experimental and theoretical approaches to conscious processing. Neuron. 2011;70:200–27.

33. Block N. What Is Wrong with the No-Report Paradigm and How to Fix It. Trends Cogn Sci. 2019;23:1003–1013. doi: 10.1016/j.tics.2019.10.001.

34. Faul F, Erdfelder E, Buchner A, Lang A-G. Statistical power analyses using G*Power 3.1: Tests for correlation and regression analyses. Behav Res Methods. 2009;41:1149–1160. doi: 10.3758/BRM.41.4.1149.

35. Kleiner M, Brainard DH, Pelli D, Ingling A, Murray R, Broussard C. What’s new in Psychtoolbox-3. Perception. 2007;36:1–16. doi: 10.1068/v070821.

36. Treder MS. MVPA-Light: A Classification and Regression Toolbox for Multi-Dimensional Data. Front Neurosci. 2020;14:289. doi: 10.3389/fnins.2020.00289.

37. Bays PM, Taylor R. A neural model of retrospective attention in visual working memory. Cogn Psychol. 2018;100:43–52. doi: 10.1016/j.cogpsych.2017.12.001.

38. Dehaene S, Changeux JP, Naccache L, Sackur J, Sergent C. Conscious, preconscious, and subliminal processing: a testable taxonomy. Trends Cogn Sci. 2006;10:204–211. doi: 10.1016/j.tics.2006.03.007.

39. Haufe S, Meinecke F, Görgen K, Dähne S, Haynes JD, Blankertz B, et al. On the interpretation of weight vectors of linear models in multivariate neuroimaging. Neuroimage. 2014;87:96–110. doi: 10.1016/j.neuroimage.2013.10.067.

40. Sergent C. The offline stream of conscious representations. Philos Trans R Soc Lond B Biol Sci. 2018;373(1755):20170349. doi: 10.1098/rstb.2017.0349.

41. Sperling G. The information available in brief visual presentations. Psychol Monogr. 1960;74:1–29. doi: 10.1037/h0093759.

42. Landman R, Spekreijse H, Lamme VAF. Large capacity storage of integrated objects before change blindness. Vision Res. 2003;43:149–164. doi: 10.1016/S0042-6989(02)00402-9.

43. Sligte IG, Scholte HS, Lamme VAF. Are there multiple visual short-term memory stores? PLoS One. 2008;3:e1699. doi: 10.1371/journal.pone.0001699.

44. Griffin IC, Nobre AC. Orienting attention to locations in internal representations. J Cogn Neurosci. 2003;15:1176–1194. doi: 10.1162/089892903322598139.

45. Souza AS, Oberauer K. In search of the focus of attention in working memory: 13 years of the retro-cue effect. Atten Percept Psychophys. 2016;78:1839–1860. doi: 10.3758/s13414-016-1108-5.

